# Diet breadth in two polyphagous *Spodoptera* moths in a wide range of host and non-host plants and the potential for range expansion

**DOI:** 10.1101/2024.07.25.605058

**Authors:** Amit Roy, Nicole Wäschke, Sophie Chattington, Roman Modlinger, Amrita Chakraborty, Thabani E.S. Chirere, Mattias C Larsson, David G Heckel, Peter Anderson, Fredrik Schlyter

## Abstract

1. Several polyphagous moths are severe crop pests. Diet breadth patterns and mechanisms among polyphagous insects provide an excellent system to study ecological and evolutionary processes in herbivores, driving dietary specialization. However, studies of diet breadth on more than a handful of crops are scarce.
2. Here, we estimated the diet breadth in two species of lepidopteran herbivores from the genus *Spodoptera*: *S. littoralis* (SL), with host range including both mono- and dicotyledonous plants and *S. frugiperda* (SF) Corn strain, primarily adapted to different grass species.
3. Larval performance on 23 crop and wild plant species from 17 families from terrestrial and wetland habitats was compared to an artificial diet in no-choice feeding bioassays. SL survived and performed better on most tested plants, particularly on the family level, except on two monocot plants (maize and leek), where SF performed well. There were five wild non-host plants where both generalists failed to survive. Nutrition indices assay corroborates the findings on a subset of plants.
4. In a subset of plants, larval feeding preference correlated partly, and larval attraction correlated well with larval performance. Female oviposition choice showed a weak correlation with larval performance. This weak correlation implies that these traits are decoupled, and other factors are crucial for female host plant selection.
5. During larval dispersal greenhouse experiments, SL and SF larvae strongly tended to migrate onto their suitable host plants, indicating that this is one factor that modulates female host plant selection.
6. In summary, SL has a broader diet breath compared to SF, surviving on wild plants with no previous exposure. The present study provides the first comprehensive data on the diet breadth of two range-expanding and highly invasive polyphagous herbivores.

## 1. Introduction

Plant-insect interaction is one of the most ancient ecological associations on earth, leading to astonishing diversity in herbivore insects. The impressive variation in resource utilization makes herbivore insects one of the most successful animal survivors in the history of life and the desired model for studying resource specialization (Futuyma & Moreno 1988). Such variations often lead to the gradient of dietary specialization among species, populations and even among individuals and mediate crucial ecological and evolutionary processes such as diversification, resistance to environmental perturbation, co-existence of competitors, etc. (Devictor, Julliard & Jiguet 2008; Büchi & Vuilleumier 2014; Hardy & Otto 2014; Forister *et al*. 2015). Microbial symbiosis and the transgenerational phenotypic plasticity of herbivore insects are often associated with fine-tuning the host acceptance and shifts (Rösvik *et al*. 2020; Roy *et al*. 2023). Studies also document that insect gene expression plasticity facilitates host shifts and aids in dealing with unexposed plant allelochemicals (Roy *et al*. 2016a; Sellamuthu *et al*. 2024). Moreover, herbivore dietary specialization or diet breadth can also be influenced by the top-down effect of the activity and efficacy of parasitoids and predators on different plants (Singer *et al*. 2014). Variations in host use may also lead to species diversification, such as in butterflies (Braga *et al*. 2018; Allio *et al*. 2021). Nevertheless, at the ecosystem level, the specialization of resource utilization can stabilize the entire ecological network of diversely interacting species (Mougi & Kondoh 2012).

By definition, the diet breadth of herbivore insects can be measured as the number of plant families they can feed on (Hardy *et al*. 2020). It is known that herbivore insect fitness profoundly relies on selecting an appropriate host. Understanding the basis underlying the host plant selection will aid in answering a key evolutionary ecological question, such as why some insects became major pests after shifting to cultivated plants from wild plants. Gripenberg *et al*. (2010) found that the female preference for optimal host plants is much stronger in oligophagous than in polyphagous insects, indicating the reduction of selection pressure due to broader host repertoire. Interestingly, the observed effect sizes for monophagous and polyphagous insects are much lower than oligophagous insects, suggesting that the preference-performance correlation is strongest among insects with an intermediate diet breadth.

With the change in global climatic conditions and rise in mean temperature, herbivore insect invasion rate also rises (Ju *et al*. 2015; Liu *et al*. 2024). Interestingly, the wider diet breadth facilitates the geographic range expansion of many herbivorous insects (Stastny *et al*. 2006; Slove & Janz 2011). While colonizing new areas, a polyphagous insect has more possibilities of finding a potential host that is closely related to its preferred host (Scriber & Ording 2005). Finding a new host in an unexplored territory perhaps eases the range expansion further beyond the original geographic range of the host plant (Gutiérrez & Thomas 2000). A broad dietary spectrum is frequently observed within invading species as they may adapt well to different and novel conditions (Goulson 2003; Granot, Shenkar & Belmaker 2017). Quantifying the diet breadth of herbivore insects will often improve the estimation of their invasiveness potential. Even if diet breadth is an essential trait, an invasive insect population cannot only be characterized by a broader diet as it is not the only precondition for becoming a successful invader (Vázquez 2006; Courant *et al*. 2017).

Recently, it has been proposed that the plant genotype and defence compounds can differentially affect the performance of insect herbivores with varied diet breadth (Damestoy *et al*. 2019; Wang *et al*. 2022). From an agricultural perspective, generalists or insects with broader diet breadth are more prone to evolve insecticide resistance or tolerance as they are already pre-adapted to a broad array of diverse phytochemicals (Dermauw *et al*. 2013; Hardy *et al*. 2018). Undoubtedly, diet breadth is one of the critical factors which influences the dynamics of herbivory and several other facets of herbivore ecology, including gut microbiome (Hardy *et al*. 2020; Šigutová *et al*. 2024; Ugwu, Wenzi & Asiegbu 2024). However, whether diet breadth affects the host shift to novel crop plants or is reflected in their host range of wild plants is unexplored. It is suggested that diet breadth is a promising avenue for future research to understand the variation in the strength of preference–performance correlation in insect herbivores (Gripenberg *et al*. 2010). Hence, an in-depth understanding of herbivore diet breadth for economically important pest insects is crucial for formulating a sustainable strategy to protect the crops. However, studies of herbivore pest insect diet breadth on more than a handful of crops are scarce.

In the present study, we evaluate and compare the diet breadth of two closely related important, invasive, and polyphagous pests: *Spodoptera littoralis* (SL, Egyptian cotton leafworm) and *Spodoptera frugiperda* (SF, Fall armyworm). Both insects are affiliated to the family Noctuidae within the order Lepidoptera. SL is one of the most destructive polyphagous agricultural pests in the subtropical and tropical range. It is expanding its geographic range with migratory populations approaching through southern Europe (i.e., eastern Spain, southern France, and northern Italy) and causing considerable damage to field crops (https://www.Cabi.org/ISC/datasheet/51070). Alternatively, SF is currently a substantial threat to Africa as its spread has continued to over 44 countries in Africa now (Ramasamy, Das & Ramesh 2022; Dessie *et al*. 2024) and even reached the Indian Ocean islands, including Madagascar (Day *et al*. 2017). It has also recently become a major pest in southern and eastern Asia, e.g., India and China (Mendesil *et al*. 2023). Apart from their economic importance, these two pests share a common evolutionary history and ecological features yet differ in their relative diet specialization scale, making them suitable for the proposed study.

We performed a series of experiments under controlled laboratory conditions to understand and compare their feeding ecology, diet breadth, and performance-preference correlation. A major no-choice feeding bioassay with 22 plants and diet control allowed covering an origin of wide systematic, cultivation, and habitat breadth, estimating larval survival and performance from neonates to adults. Subsequently, based on these results, we selected five plants; out of them, two affiliated with monocots (Maize and Leek), two are dicots (Cotton and Cabbage), and one true non-host (Malabar nut) for further downstream experiments, such as larval olfactory preference, larval feeding preference, nutrition indices analysis and female oviposition preference. Our findings fill the existing knowledge gap on the feeding ecology of these two economically important invasives and their potential for plant colonization and range expansion in new habitats under global climate change scenarios.

## 2 Materials and methods

### 2.1 Plants and insects

A selection of the experimental plants such as *Zea mays* (Maize), *Gossypium hirsutum* (Cotton), *Brassica oleracea* (Cabbage), *Ricinus communis* (Castor oil plant/ Ricin), *Trifolium alexandrium* (Egyptian clover), *Vigna unguiculata* (Cowpea), *Solanum lycopersicum* (Tomato), *Capsicum annuum* (Peppers), *Helianthus annuus* (Sunflower), *Beta vulgaris* (Beet), *Allium ampeloprasum* (Leek), *Brassica nigra* (Mustard) was grown in a Biotron chamber on a commercial substrate (Kronmull, Weibull Trädgård AB, Hammenhög, Sweden) at 23 ± 2°C and 70 ± 5% RH with 16:8 h L:D cycle until used for experiments. Other experimental plants *Mentha aquatic* (Mint), *Alisma plantago-aquatic* (Mad dog weed), *Lycopus europaeus* (Gypsywort), *Filipendula ulmaria* (Meadowsweet), *Antirrhinum austrate* (snapdragon), *Iris spuria* (Blue Iris), *Caltha palustris* (Marsh marigold), *Scutellaria galericulata* (Common skullcap) and *Phragmites australis* (Common reed) were obtained from commercial vendors (Veg tech AB, Fagerås, Sweden) and maintained under controlled environment in the greenhouse same as above. *Justicia adhatoda* L. (syn. *Adhatoda vesica* Nees) (Malabar nut) was grown in the greenhouse from the small branches obtained from Egypt. *Magnolia liliiflora* (Korean mountain magnolia) plant leaves were collected from the local gardens in Alnarp, Sweden. Detailed scientific information for the plants used in the study is given in **Suppl Appendix 1**.

The SF (maize strain) was derived from 300 larvae collected in April 2010 from two maize fields in Santa Isabel, Puerto Rico, and maintained at the Department of Entomology, Max Planck Institute for Chemical Ecology, Jena, Germany (Roy *et al*. 2016a). SL was acquired from a laboratory culture in Alnarp, Sweden, from the wild collection (2008) from Egypt. The culture is maintained on a potato-based semiartificial diet and refreshed at least once yearly with newly collected wild moths from Egypt (Zakir et al. 2013). For the present study, SF and SL were reared on a pinto bean-based semi-artificial diet for three generations at 26°C and 16:8 h L:D cycle (Perkins 1973) to maintain uniformity in the experimental setup and reduce background noise to simplify the comparisons.

### 2.2 Larval survival, performance and feeding: no-choice bioassay

Freshly hatched neonate larvae from SL and SF (maize strain) were randomly selected with a soft, thin brush and transferred in 30 ml cups containing pinto diet or pieces of fresh leaves. Each cup had a moistened plaster layer on the bottom to prevent the leaves from wilting and the larvae from desiccating (Roy *et al*. 2016a; Roy *et al*. 2016b). A single larva was placed in each cup to avoid cannibalism. Leaves or artificial diets were replaced with new ones every day. Larval survival, larval mass (10th, 14th, and 18th day), pupal mass, and the developmental time (rate) at larval and pupal stages were recorded for 50 larvae per treatment. The experiment was terminated when all larvae emerged after pupation or died. All the bioassay experiments were performed in batches with cohorts of insects on different days to minimize the plant, day, and batch (larvae) effect. Bioassay data were evaluated using IBM SPSS 22 using its descriptive routines, PCA (CATPCA) and a full factorial GzLM by GENLIN for analysis at α = 0.05 as documented previously (Roy *et al*. 2016a).

### 2.3 Host, non-host plant diet assimilation potential

Four host plants, two monocots (maize, leek) and two dicots (cotton, cabbage), one non-host (Malabar nut), and an artificial diet were used to evaluate the nutritional indices parameters in SL and SF. Newly moulted 4th instar larvae (N= 20) that had been reared on the pinto bean diet were randomly picked and starved for 24 h before they were transferred to individual cups with ca. 1000mg (fresh weight) of either plant leaf material or artificial diet). After 48h feeding, the larvae, frass, and remaining food materials were separated and dried at 60°C for 48h. Larvae were frozen for 30 min at -20 °C before drying. Initial dry weight estimations were performed by linear regression analysis for each of the diets (R2 =0.98; n=20), plant leaves (R2 =0.77- 0.85; n=20) and larval species (R2 =0.95-0.96; n=25) before and after drying in an oven. Nutritional index values were calculated as detailed by Ahn, Badenes-Pérez and Heckel (2011) and shown in the **Suppl. Tab. 4**.

Because the data of the nutrition assay showed log-normal distribution, it was transformed using logarithm base 10 before statistical analysis. Due to the presence of zeros and negative values (in all cases on Malabar nut plants), the data were transformed according to Quinn and Keough (2002), using log (Y + c +1), where Y was the variable and c was the lowest value of the variable Y. GLM regression model was used, and all interactions between factors were included in the initial model. The significance of model factors was determined by analysis of variance using a conservative F-test at the usual levels of significance: p < 0.05, p < 0.01, and p < 0.001. For the statistically significant variable, factor levels were also compared against the artificial diet using “treatment” contrasts (Crawley 2013; Pekár & Brabec 2016). All analyses were performed in the R 3.5.2 environment.

### 2.4 Preference for selected plants under multiple-choice

#### 2.4.1 Larval multiple-choice and feeding

To compare the host feeding preferences between SF and SL, feeding choice experiments were carried out with access to five different plant leaf discs (Hagenbucher *et al*. 2016). The selection of plants was the same as the nutrition indices assay. Petri dishes (diameter 14cm) with moistened filter paper (Munktell filter AB) were used to maintain the humidity and freshness of the leaves. Fresh leaf discs (∅ 2cm) were immediately placed equidistant from each other around the edge of the Petri dishes. The placement of the leaf disc was random to nullify any positional effect. A third instar larva was placed at the centre of each petri dish that was then closed with a lid and kept at 25°C, 70%RH, 16:8 L:D for 24 hours. In total, 120 replications were done for each insect species (40 replicates/day). In this setup, larvae were allowed to feed from multiple plants according to their choice. A single-leaf disc provides *ad libitum* food for third-instar larvae for 24 hours.

After 24 hours, larvae were removed from the Petri dishes. Except those of leek, the remains of all eaten leaf discs were carefully removed from the Petri dishes and taped on A4 plain copy sheets using transparent sticky tape and scanned using a printer (RICOH MPC, Sweden) at 600 dpi. The images were saved as JPEG. Due to the thickness of the leek leaves, the remains of these leaves were put on a background light projector (KAISER Prolite basic, Germany), and photos of each were taken using a Sony 12.1-megapixel digital camera. A 2cm line was drawn next to each photo/scanned image to facilitate equal image calibration during image analysis with ImageJ (http://imagej.en.softonic.com/), measuring the area consumed from each leaf disc. To calculate the dry weight per unit leaf area, 20 fresh leaves from each plant species used in the experiment were weighed using a high-precision analytical balance (HR-200, precision 0.1 mg; A&D Company, Ltd., Tokyo, Japan). The leaves were dried using a 50° C oven and weighed again to determine the average dry weight of each leaf. The total dry weight consumed was calculated using the following formulae,

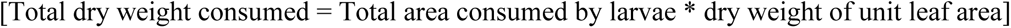

For statistical analysis, a mixed effect model with Gamma distribution was used as changes in the dry weight mass of the plant were directly correlated with the time of larval feeding. Gamma distribution requires values above zero; hence, a small constant (half of the lowest observation) was added to each value during the analysis (Pekár & Brabec 2016). We also considered the correlation between the plant selection on the Petri dishes and individual variability of the larvae nutrition intake during analysis. The correlation coefficient matrix was used in the model using an exchangeable function to eliminate the residual correlation in specific dishes.

#### 2.4.2 Larval olfactory preference

Olfactory preference was tested for SL and SF larvae between two plants using a Y-tube olfactometer (Carroll *et al*. 2006; Hagenbucher *et al*. 2016). An aquarium pump-driven flow of 0.4 l min^-1^ charcoal-filtered and moistened air was split into two separate tubes, where each tube entered a polyethylene bag containing one undamaged plant. The air from one bag was then attached to one of the test arms, and the flow from the other bag was attached to the second test arm of the sterilized Y-tube. Flow meters were used to control the airflow so that it was the same in each of the two arms.

The larvae were starved for 3-5 hours before the experiment. The olfactory preference of 4th instar larvae of SL and SF was tested, and then they were released individually into the main arm of the Y-tube. The tests were done under red light, and plants vs pure airflow (blank) was used to evaluate the experimental setup. Each larva was monitored as it moved through the main arm and into the side (test or olfactometer) arms, and a choice was recorded when the larvae had reached 5 cm into one of the test arms. The threshold time was set at >2 to <10 minutes for a given choice to be valid. Insects that failed to choose within the given time were excluded from the analysis. Each larva was only used once. All possible combinations between cotton, cabbage, and maize plants were tested multiple times with small groups of larvae on different days to minimize the plant, day, and larval batch effects. Plant positions were also randomized, and the olfactometer was cleaned after each set of experiments.

#### 2.4.3 Larval migration between plants

The dispersal behaviour of SL and SF larvae was measured when a choice between maize (monocot) and cotton (dicot) was provided (Anderson, Sadek & Wäckers 2011). For the experiment, potted plants of maize and cotton were placed in pairs on plastic trays, close enough to allow contact and larval movement between their leaves. For each setup, connections between leaves of similar height were established in three places using pieces of steel wire. The pots were placed in smaller dishes, and the trays were then filled with water to approximately 2 cm height to prevent larvae from escaping. A piece of cardboard, with two holes cut where the stems of the plants, was used in every setup to cover the soil of the two pots. It provided a bridge between the plants so the larvae could migrate between plants at the soil level.

The setups were placed in a greenhouse chamber (23-27°C, 16:8 h L:D). At the start of the experiment, five 3rd instar larvae were placed on separate leaves on either the cotton or the maize plant in each experimental setup. For the controls, five larvae were placed on separate leaves of one of two conspecific plants. The setups were checked every 8 or 16 hours, and the number of larvae found on each plant was counted for 72 hours. Identical experimental replicates were made multiple times (n=15) with small groups of larvae on different days to eliminate the plant, day and larval batch effect. To evaluate the migration of SF and SL larvae, 40- and 72-hour periods were chosen (Anderson et al. 2011), and the number of larvae present on the different plants was compared. Movement from cotton to maize or maize to cotton was tested separately by Poisson GLM and χ2 test.

#### 2.4.4 Oviposition choice

The oviposition choice of SL (preferring dicots) and SF (preferring monocots) was tested using undamaged plants of the two classes. The experiments were performed in a four-choice situation by offering the dicots cotton, cabbage, and monocots leek and maize as oviposition plants (Hagenbucher *et al*. 2016). Plastic cages (30 x 30 x 30 cm BugDorm-1, Mega View Science, Taichung, Taiwan) with sidewalls of nylon tissue (mesh size ca. 1 mm) were placed in a climate chamber (25 ± 2°C, 70 ± 2% RH, and 16:8h L:D). Freshly cut mature leaves of similar fresh weight of the four plant species were inserted singly into plastic tubes (∅ 2.5 cm, 8 cm high) filled with water and plugged with a cotton ball. Assays using cut leaves have previously been shown to give similar results compared to using entire intact plants (Thöming et al. 2013). To avoid positional effects, the positions of the plant species were randomized. Leaves from the same plant were used for two cages. Two to three-day-old adults were allowed to mate. Copulating moths were introduced in each cage, and females were allowed to oviposit for two nights. The egg batches were counted, collected on day three, and weighed using an analytical balance (HR-200, precision 0.1 mg; A&D Company, Ltd., Tokyo, Japan). The number of eggs was calculated using the equation [Number of eggs = weight of eggs (mg) x 20] (Zakir *et al*. 2013). To compare the oviposition preference of the females for the four different plant species, we use non-parametric statistics, overall Friedman ANOVA, followed by Wilcoxon signed-rank test suitable for non-homogenous data set. P-values were adjusted by the "Holm" method (Haynes 2013). Data were analyzed using R (Version 0.98.1091, 2014).

## 3. Results

### 3.1 Larval survival and performance under no-choice at a wide array of host and non-host plant species

*At the level of plant species*, SL overall, as the broader generalist among the two, survived and performed better on the leaves of most of the tested plants than the SF (maize strain). On the semi-artificial pinto bean diet, the response of the two insect taxa was equivalent, as reported earlier (Roy *et al*. 2016a). However, dramatic effects of host/non-host plant feeding were observed for larval survival and performance between and within taxa (**Fig. 1**, Plant species ordered by increasing survival).

**Fig. 1:**
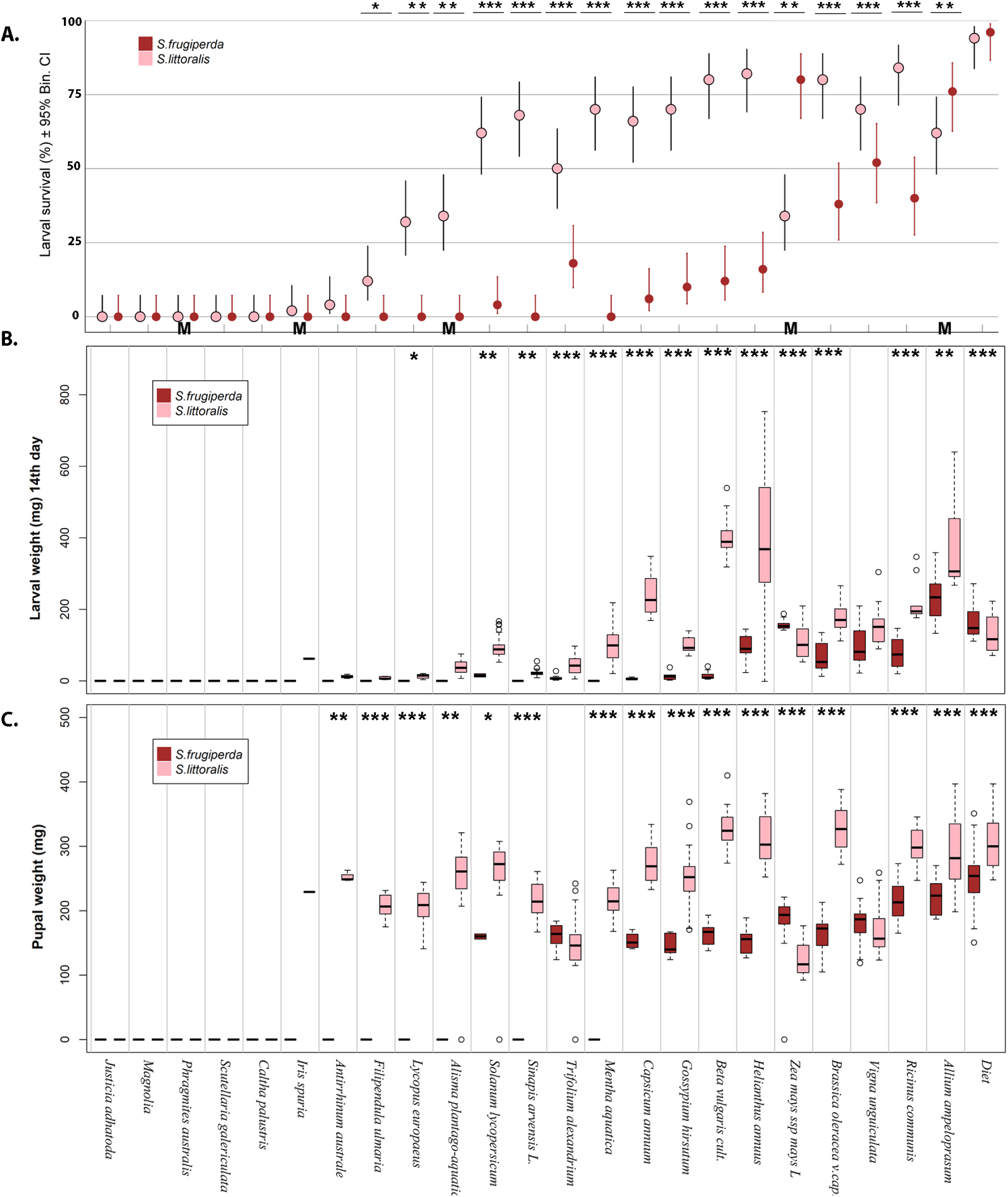
No-choice test on replenished leaf disc in Petri discs for total 23 plants. **A**) Larval survival of SL and SF on the diverse array of host and non-host plants and artificial diet with exact binomial confidence levels (Newcombe 1998; Newcombe 2012; Fagerland, Lydersen & Laake 2015), **B** & **C**) Larval weight (14th day) and pupal weight of SL and SF after eating the 24 foods (medians, interquartile range, extremes and outliers plotted). Analysis: One-way ANOVA compares the performance difference between SL and SF on a particular host plant. Survival compared confidence intervals overlaps or not. **M** = Monocots. *) *p* < 0.05, ** *p* < 0.01, and *** < 0.001

#### 3.1.1 Survival

Survival on the semi-artificial diet was high for both insects, ≈ 95%. On the 23 plant leaves, the survival pattern between the two insects is dramatically different, with SL showing overall higher survival with some survival on 18 of 23 plant species and diet compared to only 11 for SF (**Fig. 1A**). Overall, the SL survival is higher, 41.8 ±6.8% (mean± SE, n= 23) compared to 15.3 ±5.1% for SF (species different at Z= −3.07, Asymp Sig *p* =0.002 by Wilcoxon S-R). The effect size is large, and its confidence interval does not include zero (Hedges’ *g ±*95% CI: 0.90 ±0.61)(Nakagawa & Cuthill 2007). The SF response had a high relative variability (coefficient of variation= 156%) compared to SL (76%). More than >70% of larval survival was observed when the larvae (SF) were fed on plant leaves from the two monocots, maize (*Zea mays*) and leek (*Allium ampeloprasum*) (**Fig. 1A**). There was no survival of larvae for five plant taxa for any insect; three of these taxa came from wetland habitats.

For SF, 12 plants gave zero survival; of these plants, eight were aquatic (**Suppl. Appendix 1**). In 12 plants, SL had significantly higher survival, while SF was a better survivor only on one monocotyledon plant (maize) (**Fig. 1A**, monocot plants marked **M**). SF larval survival on cowpea was around 60%, whereas on castor oil plants (*Ricinus communis*), cabbage (*Brassica oleracea*), sunflower (*Helianthus annuus*), beet (*Beta vulgaris*), and Egyptian clover (*Trifolium alexandrium*) leaves were 40-20%. Approximately ≤10% SF larval survival was observed on cotton (*Gossypium hirsutum*), mustard (*Sinapis arvensis*), tomato (*Solanum lycopersicum*), and Paprika (*Capsicum annuum*) leaves.

SL survival was more than 80% on castor oil plants, sunflowers, and beet leaves. Larval survival ranged from 50-80% on cabbage, cowpea (*Vigna unguiculata*), cotton, mustard, peppers, water mint (*Mentha aquatica*), tomato, and leek leaves. Larval survival <50% was observed when SL larvae were fed on maize, mad-dog weed (*Alisma plantago-aquatica*), Egyptian clover, gypsywort (*Lycopus europaeus*), meadowsweet (*Filipendula ulmaria*), while only <10% on snapdragon (*Antirrhinum austrate*), blue Iris (*Iris spuria*), Malabar nut (*Justicia adhatoda*), marsh marigold (*Caltha palustris*), common skullcap (*Scutellaria galericulata*) and common reed (*Phragmites australis*) leaves (**Fig. 1A**).

#### 3.1.2 Performance

Due to mortality on many sub-optimum diets (i.e. non-host plant leaves), the original 2240 cases were reduced to 840 cases of larval plant-feeding (937 including the semi-artificial diet). The dependent variables were, as expected, well correlated (**Suppl. Tab. 1**). In a simple PCA of two dimensions, with the nine dependent performance variables (body masses, etc.) and the nine independent factors (insect, plant, and ecological and systematic levels), the larval mass at 14 days showed the highest loading among the dependent variables (*i.e.* was the most important dependent in separating the plant/insect combinations) (**Suppl. Fig. 1**). Several dependents, such as pupal weight and days to pupation, had similar loadings (**Suppl. Tab. 2**). Larval mass at 10 and 18 days, respectively, gave less clear patterns than at 14 days (**Suppl. Fig. 2A, B**). Performance, as larval mass at day 14, varied widely with insects and plants (**Fig. 1B**), but SL larvae were at the top for almost all plants, while the two insects were similar when reared on the semi-synthetic diet. Overall, SL larval mass was higher 193 ± 11.0 mg (mean ± SE), 651 (n) compared to SF 121 ± 6.8, 286. The effect size is medium, but its confidence interval does not include zero (Hedges’ *g ±*95%CI: 0.30 ±0.14).

Pupal weight shows many significant differences between the two moths; in all but five plant species, they are lower in SF than in SL, but were more similar for the semi-synthetic diet (**Fig. 1C**). Days for pupation have narrower distribution than mass, with both median and minimum at ca 20 days for suitable plants and maxima at ca 40 for both insects (**Suppl. Fig. 2C).**

The *Insect* (SL and SF) and *Plant* factors (23 species, the lowest plant systematic level) in two-way ANOVA on larval mass at 14 days had both highly significant influences alone (*p* <0.001, **Tab. 1A**), as had their interaction (*p* <0.001). *At the level of Division* (Monocots /Eudicots), this highest plant taxon level did not affect feeding success alone (Wald χ1^2^= 1.03, *p*= 0.31) by a two-way GzLM for larval mass at 14 days (Deviance 1142, df 837, Value/df 1.4; Likelihood Ratio χ^2^ 45.6, df 3, *p* <0.0001) (Analysis same type as in **Tab. 1A**). However, there was a strong effect of factor Insect alone (Wald χ1^2^= 32.8, *p*= <0.0001), but no sign of any interaction of factors Division × Insect (Wald χ1^2^= 0.06, *p*= 0.80).

**Table 1.**
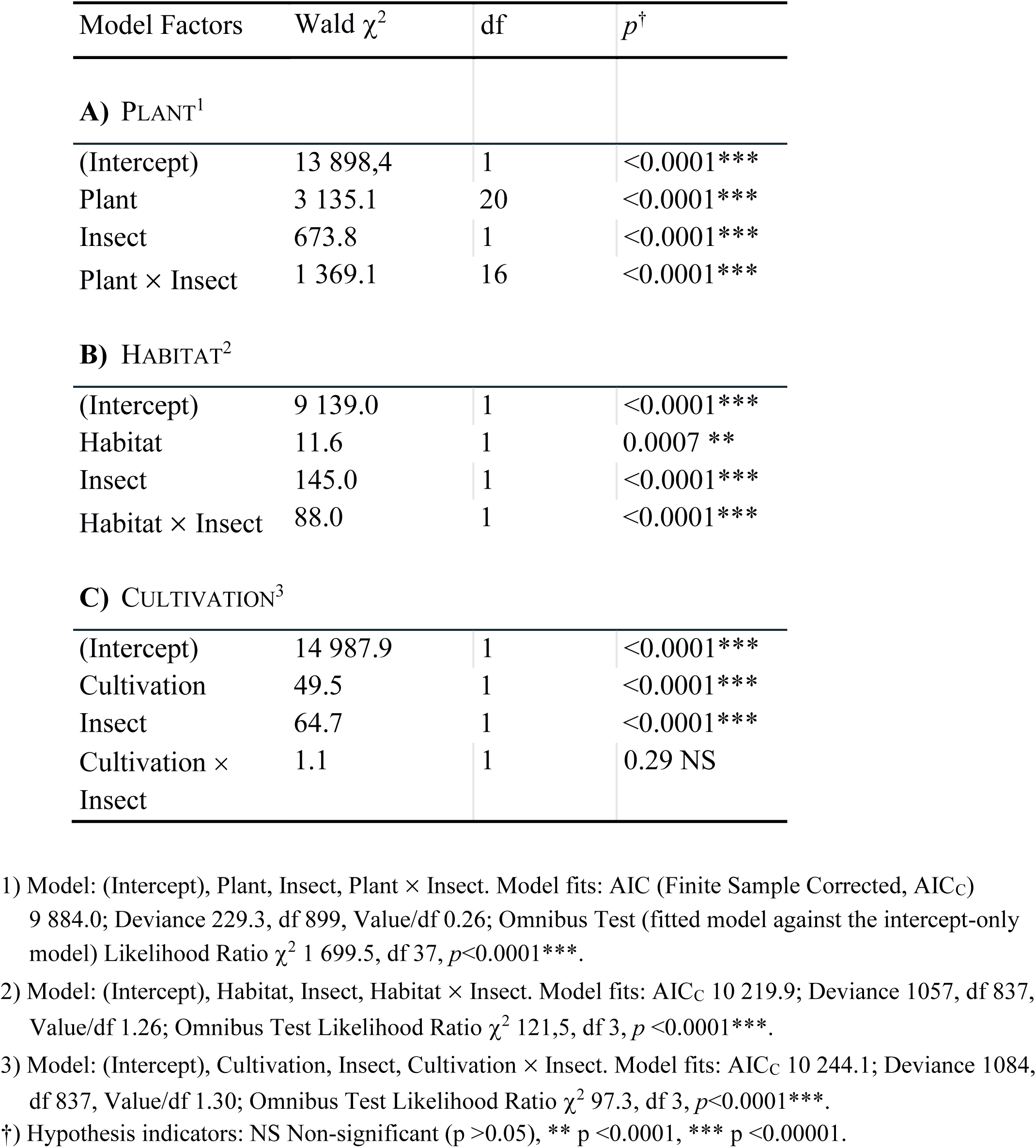
GzLM model effects type III for insect larval weight on day 14 by plants, habitats, and cultivation status with gamma distribution and log link.

*At the level of plant Families*, the non-significant effect of the highest taxon level (Division) is well illustrated by the considerable overall variation of performance at all the lower taxa levels: Class, Order, and down to Family level, visible for larval mass at 14 days (LW14) in **Tab. 2**.

**Table 2.**
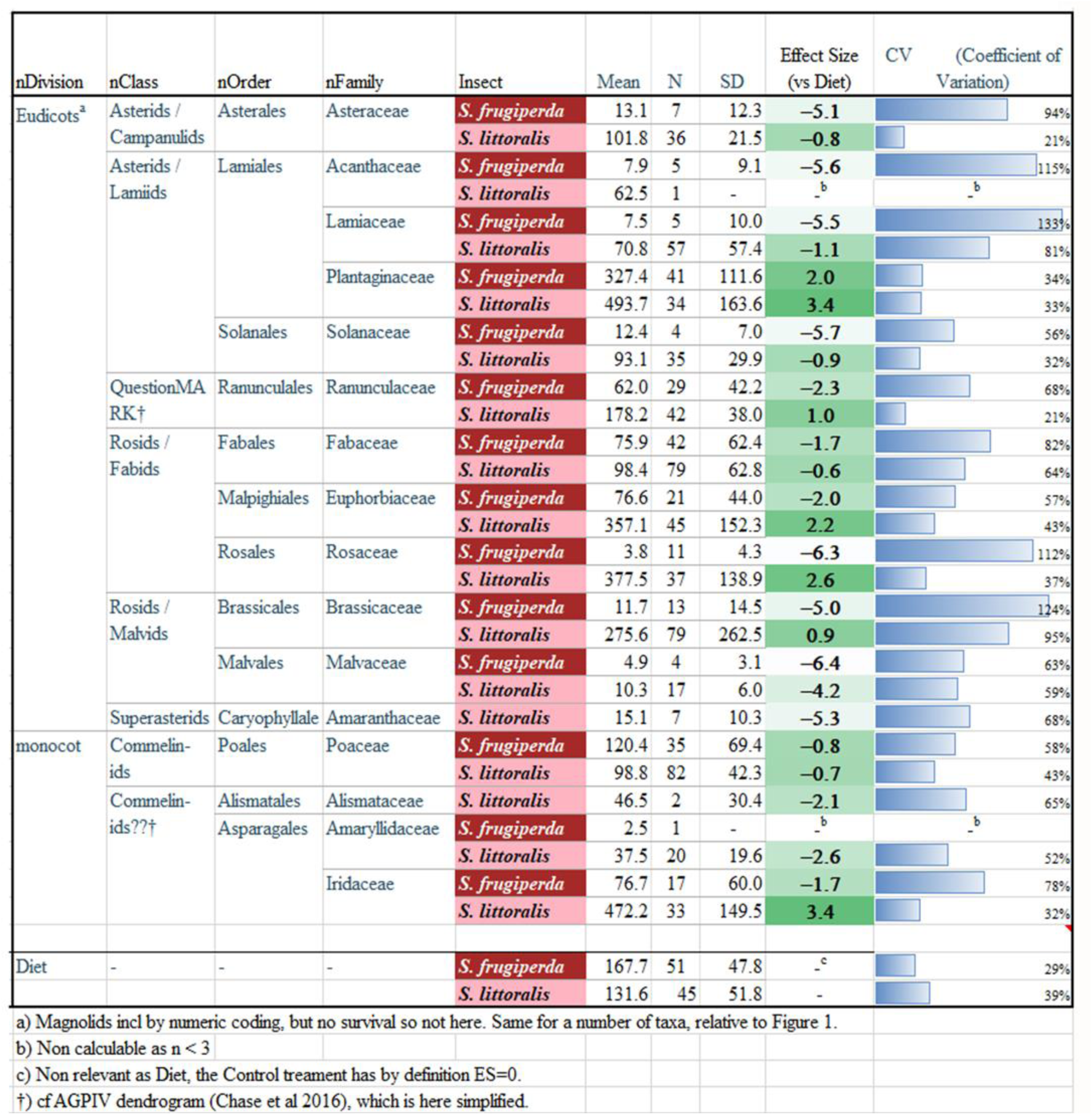
Feeding performance by nested plant taxon levels (Chase et al., 2016) compared to diet: Larval mass at 14th-day means and SD of estimates for the two insects, with effect size and relative variation.

Not only are the mean performances as LW14 different between the two insect species at plant Family levels compared to diet. The effect sizes for LW14 vs diet for the two moths ranged widely (*g=* −6.4 to +3.4 units, 7 cases above zero, 21 cases below zero) (**Tab. 2**, second to last column). In addition, the relative variation within insects is high (21 -133 %CV) but low for the artificial diet (≈ 30%) and more strongly so for families with a low average performance by an insect (**Tab. 2**, last column). The latter is particularly strong for SF data for several plant families, where its larvae could barely survive and grow.

*At nested plant taxon levels (from Division to Family):* The variation in performance by subsequent taxon levels down to family level is noticeable (**Tab. 2**). However, as the systematic levels of the plant taxa are hierarchical, the lower ones are factors not independent but are nested under the higher levels, and thus each lower level is not an independent factor level as assumed for standard analytical statistics such as ANOVA. Instead, they are biologically nested under their next higher hierarchical step (diet should not be included, as not a plant taxon). In the nested approach for the two insects, the highest level of Division (Eudicots, Monocots) alone again showed surprisingly little or no importance (Nested ANOVA **Tab. 3**, *p*= 0.37) while factor Insect alone was (*p* <0.0001). In contrast, each of the other lower-level plant factors alone, such as Order, Class, and Family, were all as nested factors highly significant (*p* <0.0001, **Tab. 3**).

**Table 3.** GLzM model^1^ effects type III for larval weight at day 14 by Insect and nested plant taxa levels with gamma distribution and log link.

| Model factors | Wald $\chi^2$ | df | $p$ † |
| --- | --- | --- | --- |
| (Intercept) | 2907.00 | 1 | <0.001*** |
| Insect | 127 | 1 | <0.001*** |
| Division | 0.82 | 1 | 0.366 NS |
| Class (Division) | 242 | 6 | <0.001*** |
| Order (Class(Division)) | 292 | 5 | <0.001*** |
| Family (Order(Class(Division))) | 372 | 3 | <0.001*** |

†) Model fits: AIC (Finite Sample Corrected, AIC<sub>C</sub>) 9 803; Deviance 824, df 824, Value/df 0.82; Omnibus Test (fitted model against the intercept-only model) Likelihood Ratio $\chi^2$ 564.2, df 16, $p < 0.0001$ \*\*\*. †) Hypothesis indicators: NS Non-significant ( $p > 0.05$ ), \*\* $p < 0.0001$ , \*\*\* $p < 0.00001$ .

*At the two ecological levels*, Habitat and Cultivation status (**Suppl. Annex 1**), the factor *Habitat* (Terrestrial/Wetland) was by itself significant (*p* = 0.0007) while *Insect* highly so (*p* <0.0001), and the interaction of *Habitat* × *Insect* was also highly so (*p* <0.0001, **Tab. 1B**) by two-way GzLM for LW14. In other words, the response to *Habitat* was not only significant between, but in opposite direction for the two insects, where the higher mass of SL was a mirror image of the much lower mass in SF when feeding on level “Wetland” plants when plotted (**Fig. 2A**).

**Fig. 2.**
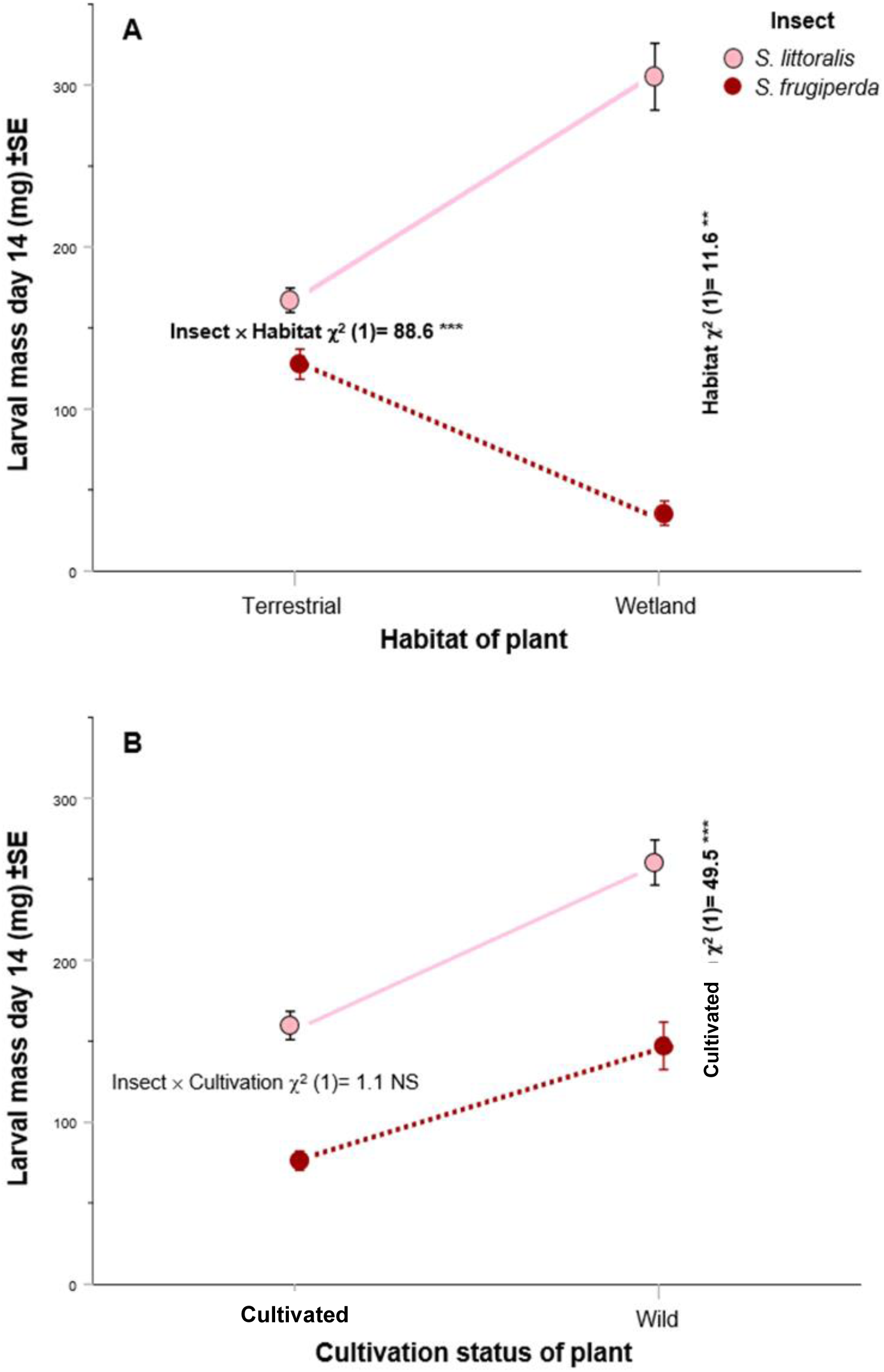
Factorial interactions of ecological factors for larval mass for the two species. **A**) Insect × habitat interaction, **B**) Insect × cultivation interaction. Significant interactions and ecological factors indicated by stars and bold text (\*\**p* < 0.01, *** *p* < 0.001, and NS Non-significant) by Generalized Linear Model GENLIN (**Tab. 1**). Factor Insect highly significant in both comparisons (*p* <0.0001, **Tab. 1 B & C**). The strong interaction in A shows clearly a different reaction norm^3^ for the two species. Data points (Means ± SEM) on untransformed scales.

The *Cultivation status* factor (Cultivated / Wild) gave a response pattern strikingly different from the habitat factor for the larval masses at 14 days. The single factors, *Cultivation status* and *Insect*, were both highly significant for feeding response as LW14 (*p* <0.0001). In contrast to Habitat factor, the interaction with factor Insect was far from significant (*Cultivation status* × *Insect*, *p* =0.29, **Tab. 1C**). In other words, *Cultivation status* level “Wild” gave a similarly larger mean for both insect species (**Fig. 2B**). Thus, reaction norms were in the same direction, but differed in magnitude between the two insects (**Fig. 2**).

#### 3.1.3 Nutritional indices of five selected plants and artificial diet

Six growth and energy conversion indices after no-choice feeding on selected plants (two monocot host plants, two dicot host plants, the artificial diet, and a non-host plant) were estimated for fourth instar larvae from SL and SF (**Fig. 3, Suppl. Tab. 3**). The comparison with performance at diet as control for the two moths show effects for all the five plants, but most strongly and similar for Malabar Nut, with effect sizes (*g*) substantially low (-8.4 to -2.7) for the six indices in the two moths. In total of 60 data point combinations 55 had effects values |*g*| >1; in other words, there were larger than 1 SD differences between Control (Diet) and Treatment (Plants) food, which is a large effect for >90% of combinations (**Suppl. Tab. 4**). Overall results from ANOVA showed a significant interaction between factors *Diet* (fed on) and *Insects* (**Suppl. Tab. 5A**), indicating the contribution of the diet (plant or artificial) to the variance between growth or energy conversion rates between SL and SF. By and large, the six indices plotted show very similar patterns over food (five plants and artificial diet) and insect combinations (**Fig. 3**), with the of fresh body mass increase showing the most substantial variation (R^2^ Adj.= 0.88, **Suppl. Tab. 5A**).

**Fig. 3.**
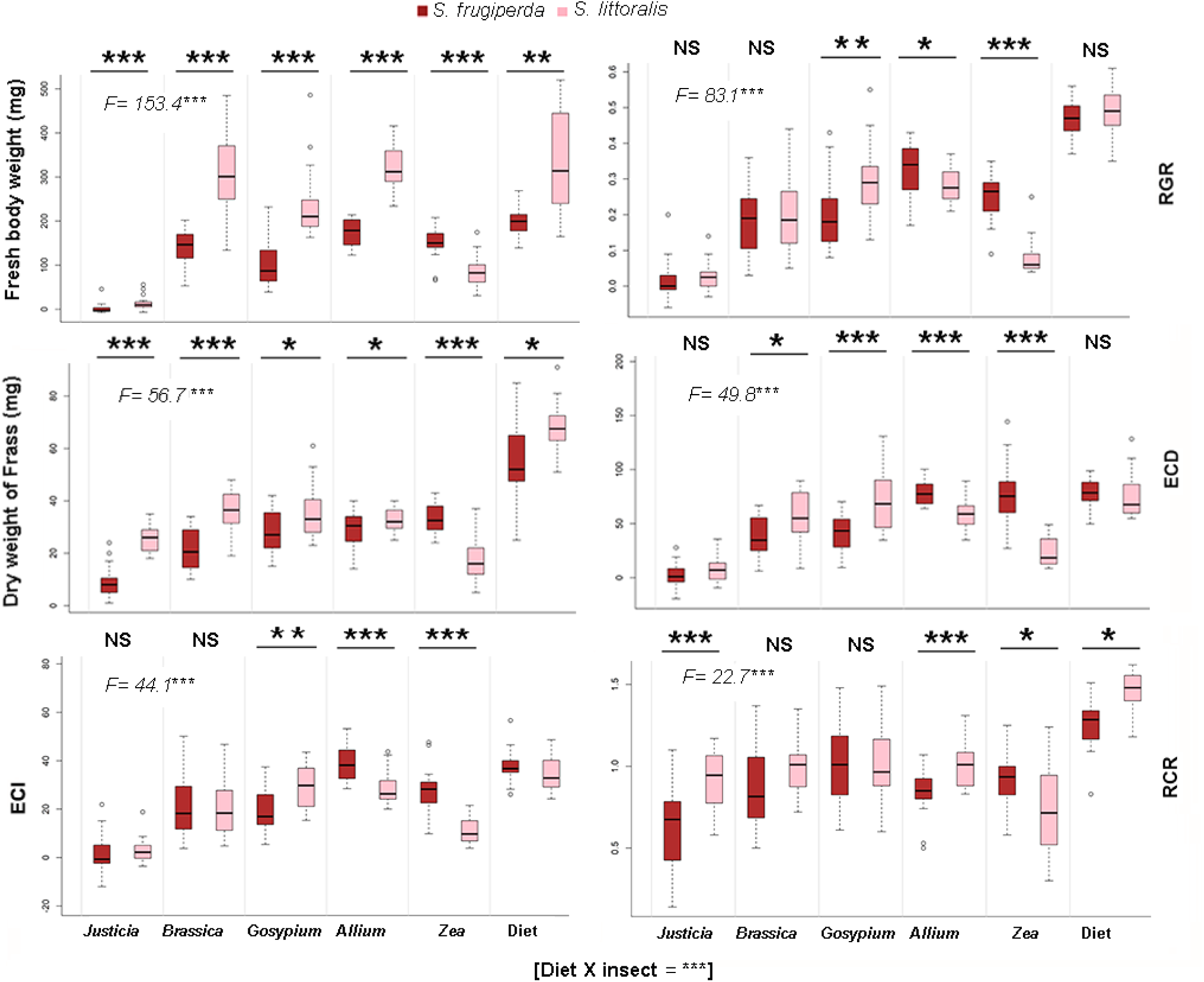
Six nutritional indices parameters for SL and SF on five different plants and the artificial diet, sorted by decreasing *F*-value (overall model GzLM, Suppl. Tab. 5A). Box plots as in Fig. 1. Acronyms: Relative growth rate (RGR), Efficiency of conversion of ingested food (ECI), Efficiency of conversion of digested food (ECD), relative consumption rate (RCR). Indices are mathematically defined in **Suppl. Tab. 3**. Statistics: Descriptive with effect size values -see **Suppl. Tab. 4**. Nutrition indices parameters were compared using five contrast matrices with different insects on the same plant. Analytical: The *p*-values for *F*-test and levels of significance are: * *p* < 0.05, ** *p* < 0.01, and *** < 0.001 (further analytical details in **Suppl. Tab. 5 B & C**).

Responses to the artificial diet by the indexes are less variable, and only a few significant differences between the two insects exist for all indices (3 of 6 possible are NS). In contrast, for responses to plant foods, there are many more significant differences; 23 of 30 possible have *p* < 0.05 or less (**Fig. 3**). Most plant-indices combinations (29 of 36) have larger values for SL median than for SF (**Fig. 3, Suppl. Tab. 5B**). The only exceptions are feeding on maize (all 6 cases) and half of those for leek, where SL has a higher value for two indices (**Suppl. Tab. 5C**).

Considering the nutritional indices parameters for non-host Malabar nut plants, both insects differ considerably in wet body weight (Bw, *p* <0.001), frass production (F, *p* <0.001), and RCR (*p* <0.001), indicating better dietary utilization of the broader generalist SL, even on a non-host plant (**Suppl. Tab. 5C**).

### 3.2 Preference behaviours for selected plants under multiple-choice

#### 3.2.1 Larval feeding preference for five selected plants

The experimental scheme of a multiple-choice assay (disc assay) feeding preference with five plants is given in **Fig. 4A**. Considering the total amount of plants consumed, SL larvae consumed two times more than SF larvae, *i.e.*, comparing the consumption of same plant between *Spodoptera* taxa, SL consumed a substantially higher amount of cotton (p <0.001), cabbage (p <0.001), leek (p <0.001) and Malabar nut (p = 0.017). In contrast, SF larvae only consumed higher amounts of maize (p <0.001, last plant in **Fig. 4B**). In descending order, most SL larvae had fed on cabbage, cotton, leek, maize, and Malabar nut (**Fig. 4C**). Investigation of larval feeding preference after 24 h confirmed that all SL and SF larvae (except one) had fed on at least one plant. More than 65% SL larvae fed on multiple plants, whereas only around 35% SF larvae fed on more than one plant, confirming SL as a broader generalist among the two, with a higher tendency to feed on all the available plants (**Fig. 4D**, **Suppl. Fig. 3**). Malabar nut was the least consumed plant species, regardless of moth species, which endorses previous findings from the nutritional indices assay.

**Fig. 4.**
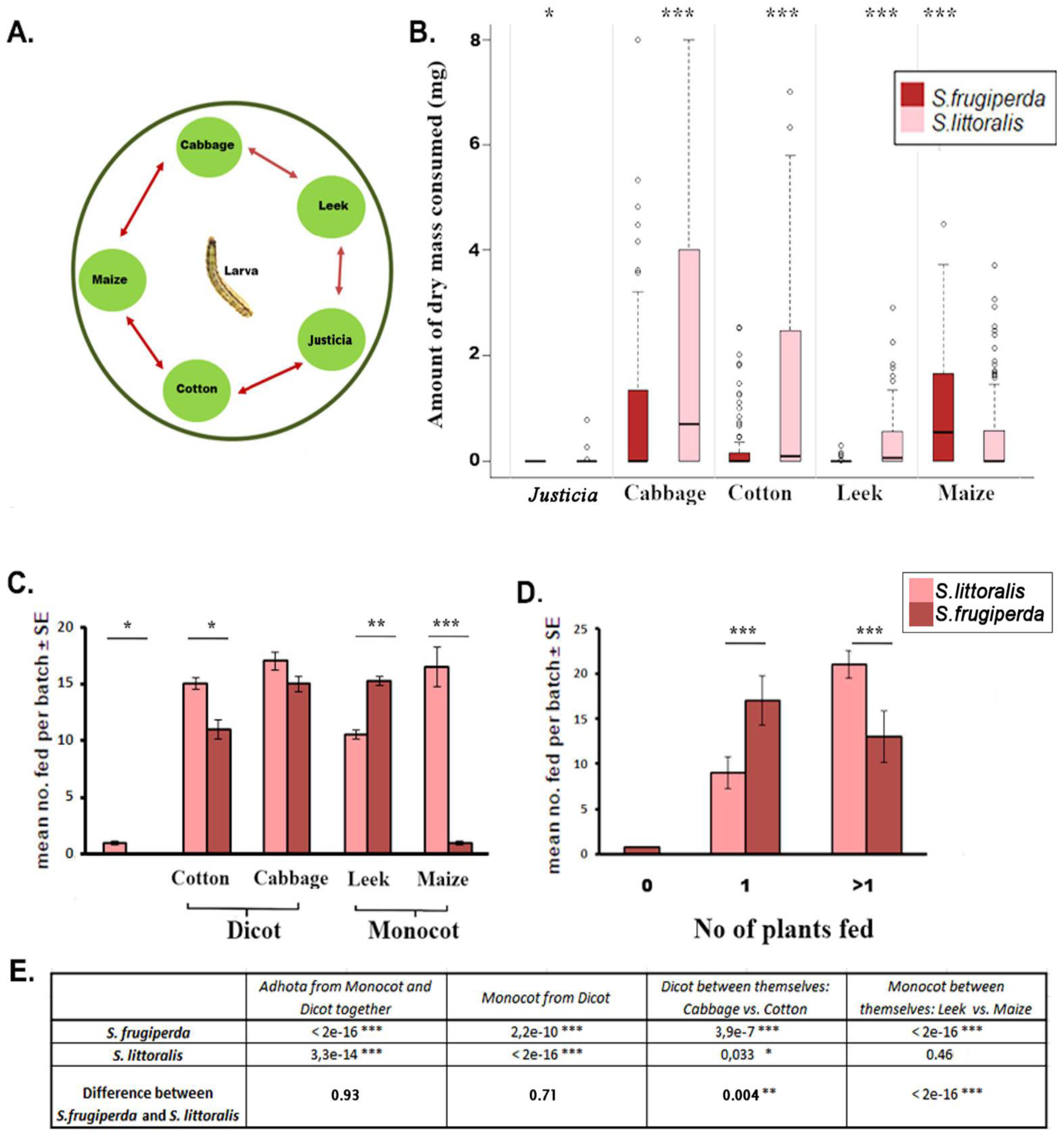
Larval feeding under multiple choice of five plants in disc assay (*n*=12 per species). **A)** Scheme of the disc assay where larvae have equal opportunity to choose the plant to feed, **B)** Amount of plant dry mass consumed by SL and SF, **C**) Mean number of larvae per batch fed on different plants after 24 hours, **D**) Number of plants SL and SF larvae fed per batch (see **Suppl. Fig. 3** for statistical details). Box plots as in Fig. 1. **E**) Contrast *t*-test using four contrast matrices based on the best GLM model. The p-values for t-test are: * *p* < 0.05, ** *p* < 0.01, and *** *p* < 0.001.

Differences between the consumption of monocots from dicots were also significant within SL (p <0.001) and SF (p <0.001) (**Fig. 4E**). The difference in dicot consumption in the SF was also vivid; cabbage was preferred more than cotton (p <0.001). Unexpectedly, SL consumed more cabbage than the presumed main host cotton (p = 0.03). Within dicot (p = 0.003) and monocot (p <0.001), there were substantial preference differences between *Spodoptera* taxa (**Fig. 4E**). While in SF, the preference for maize was entirely prevailing (p <0.001), in SL, there was no preference difference between maize and leek (p = 0.4, **Fig. 4E**).

#### 3.2.2 Larval olfactory-based preference in paired test of three plants

The Y-tube choice test of three pairs of plants demonstrates that both SL and SF had significant preferences between cotton and maize: SL preferred cotton over maize (*p*= 0.001), whereas SF displayed the opposite trend (*p* <0.001) as anticipated (**Fig. 5 A**). However, SL did not show any significant preferences in the two other choices between cotton-cabbage and cabbage- maize, but for SF, maize was significantly preferred over cabbage (*p*= 0.01) (**Fig. 5 A, C**). The GzLM binomial analysis also showed a significant effect of the interaction between factors (*Plant* × *Insects*, *p* <0.001) on larval insect olfactory-based preferences (**Fig. 5 B**). Correspondingly, comparing preference differences for each plant set at the insect taxon level, the preferences for cotton-maize (*p* <0.001) and cabbage-maize (*p*= 0.03) were considerably different between SL and SF (**Fig. 5 C**).

**Fig. 5.**
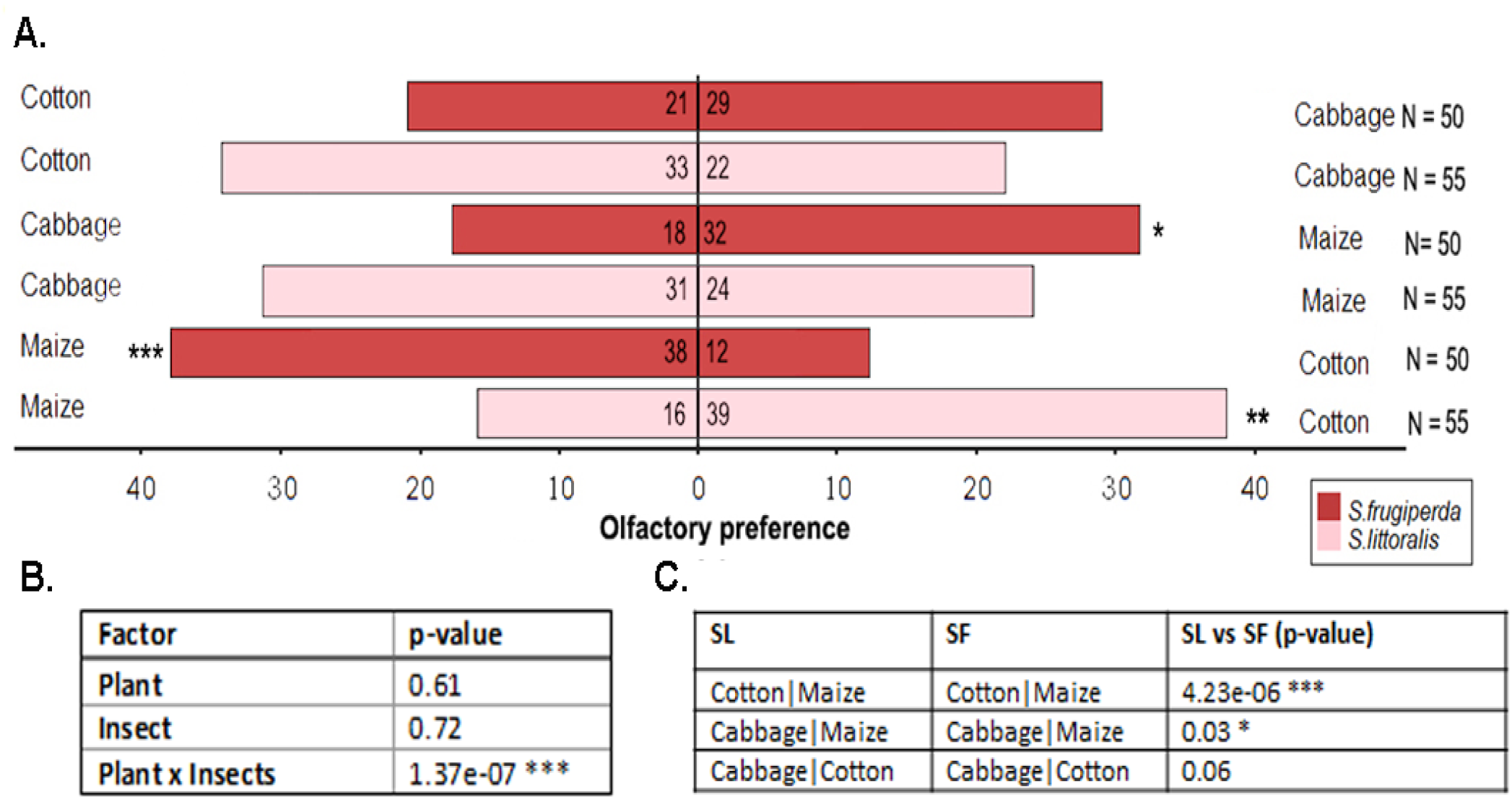
Larval olfactory preference for three plants (Cotton, Cabbage, and Maize) in a Y-tube choice assay. **A**) SL and SF larval olfactory preferences as numbers entering a Y-tube arm. The difference in the plant preference in SF and SL was evaluated using contrasts after GzLM binomial analysis. **B**) An ANOVA-like test for the interaction of factors was performed using GzLM binomial analysis. **C**) Statistical tests (using contrasts) of larval olfactory preference difference between SL and SF for all plant combinations. For each set of comparisons, approximately 15-20% of tested insects did not choose any plant within the given time, hence excluded from the analysis. Thus, N shows only those larvae that made a choice within the given time. Symbols for *p*-value levels: *) *p* <0.05, **) *p* <0.01, ***) *p* <0.001.

#### 3.2.3 Larval dispersal between primary host plants

We tested larval dispersal to assess whether SF and SL larval host plant preference translated to an increased motivation to migrate to a primary, preferred host plant when offered an adequate plant host for feeding. Dispersal behaviours for SL and SF larvae from cotton to maize or vice versa were monitored for 72 hours in the greenhouse. SF larvae, placed initially on preferred host maize, displayed little migration to cotton (**Fig. 6 A**; p < 0.001 after 40 and 72 h). The SF larva behaved in striking contrast when placed on cotton, dispersing more from cotton to maize (**Fig. 6 B**; p < 0.001). However, when SL larvae were placed initially on the non-preferred plant maize, they showed a consistent migration from maize to cotton compared to SF larvae until the end of the experiment (**Fig. 6 C**). Like SF, SL larvae migrated very little to maize when placed initially on their preferred cotton (**Fig. 6 D**). These trends were consistent with all larval counts at time points from 16 h onwards. Thus, in a visually striking mirror-like fashion, larvae moved consistently from cotton to maize or vice versa when comparing (**Fig. 6 A** and **D**) between the two *Spodoptera* spp. (**Fig. 6 B, C**). Hence, both SL and SF larvae demonstrate a similar adaptive behavioural response to finding a suitable host by olfaction over distance.

**Fig. 6.**
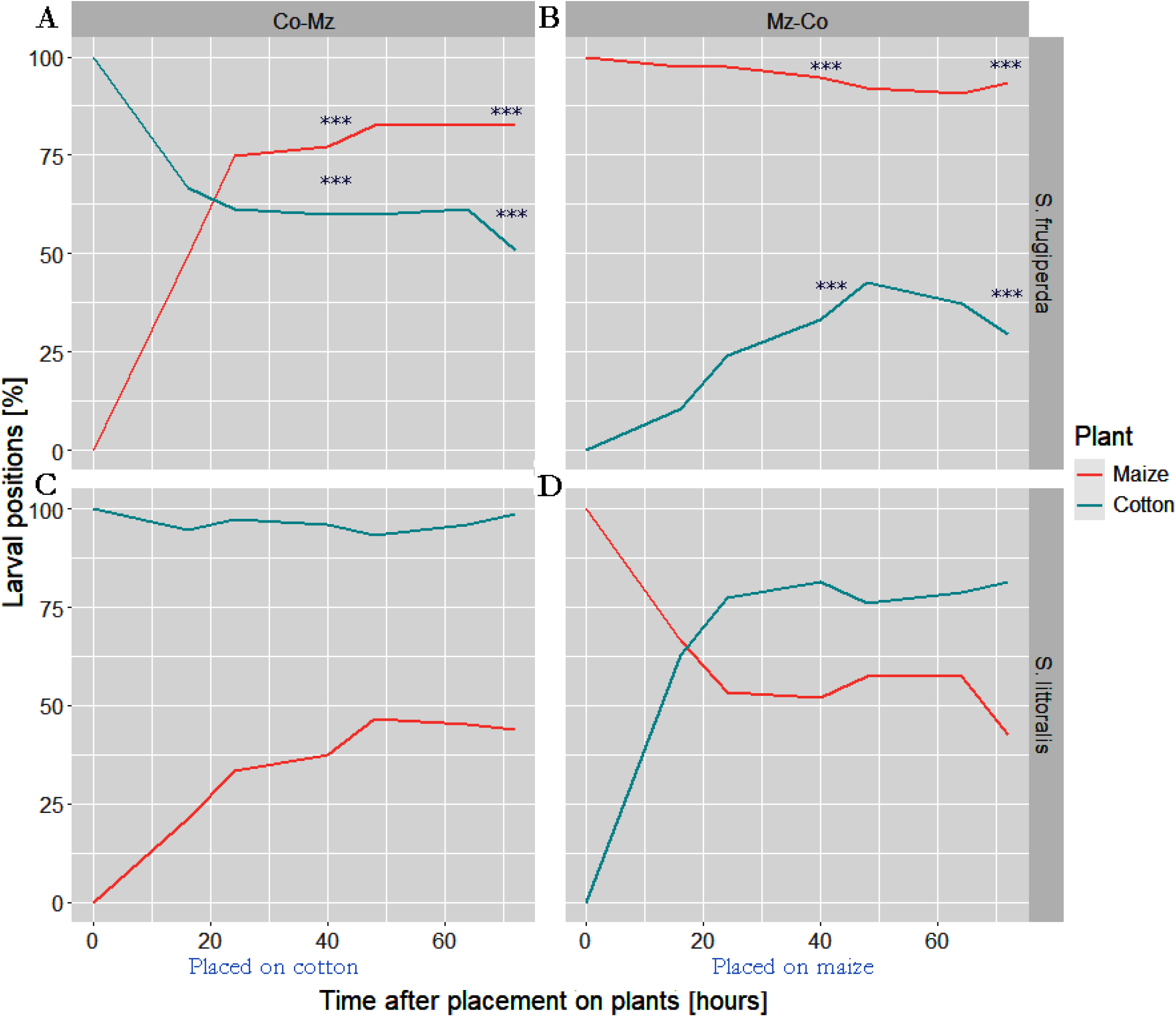
Larval dispersal between primary host plants. Larvae were allowed to disperse freely between and choose to feed on healthy cotton vs a healthy cabbage plant, their preferred host plants, respectively. In each tested pair of plants (15 pairs/combination/ species), five third instar SF or SL larvae were placed on one plant (cotton or maize). In addition, larval dispersal between plants was observed. The graphs show the cumulative number of experiments on plant pairs where the SF (**A** and **B**) or SL (**C** and **D**) larvae had either stayed at the first plant or migrated to the other plant at different observation times. In all panels, the larvae not recorded on any plant are omitted for clarity, as the number can be calculated from the total 100% and was around 20 -40%, but never different between species (*p* >> 0.05). Symbols for *p*-values of differences between species (**A** vs **C** and **B** vs **D**) of the larval positions from chi-square test for data after 40 and 72 h are: **) *p* <0.01, *** *p* < 0.001.

#### 3.2.4 Female oviposition choice among four selected plants

Comparing oviposition site preference for SL and SF, we allowed mated females to choose between four plant species (cotton, cabbage, maize, and leek) (**Fig. 7 A**, insert). The average egg mass per SL mated female was highest on cotton followed by maize, but few on cabbage or leek, whereas in the SF females, the highest amount of egg mass was found on maize followed by cotton, and again few on cabbage or leek (**Fig. 7 A**). The number of eggs followed the same pattern (**Fig. 7 B**).

**Fig. 7.**
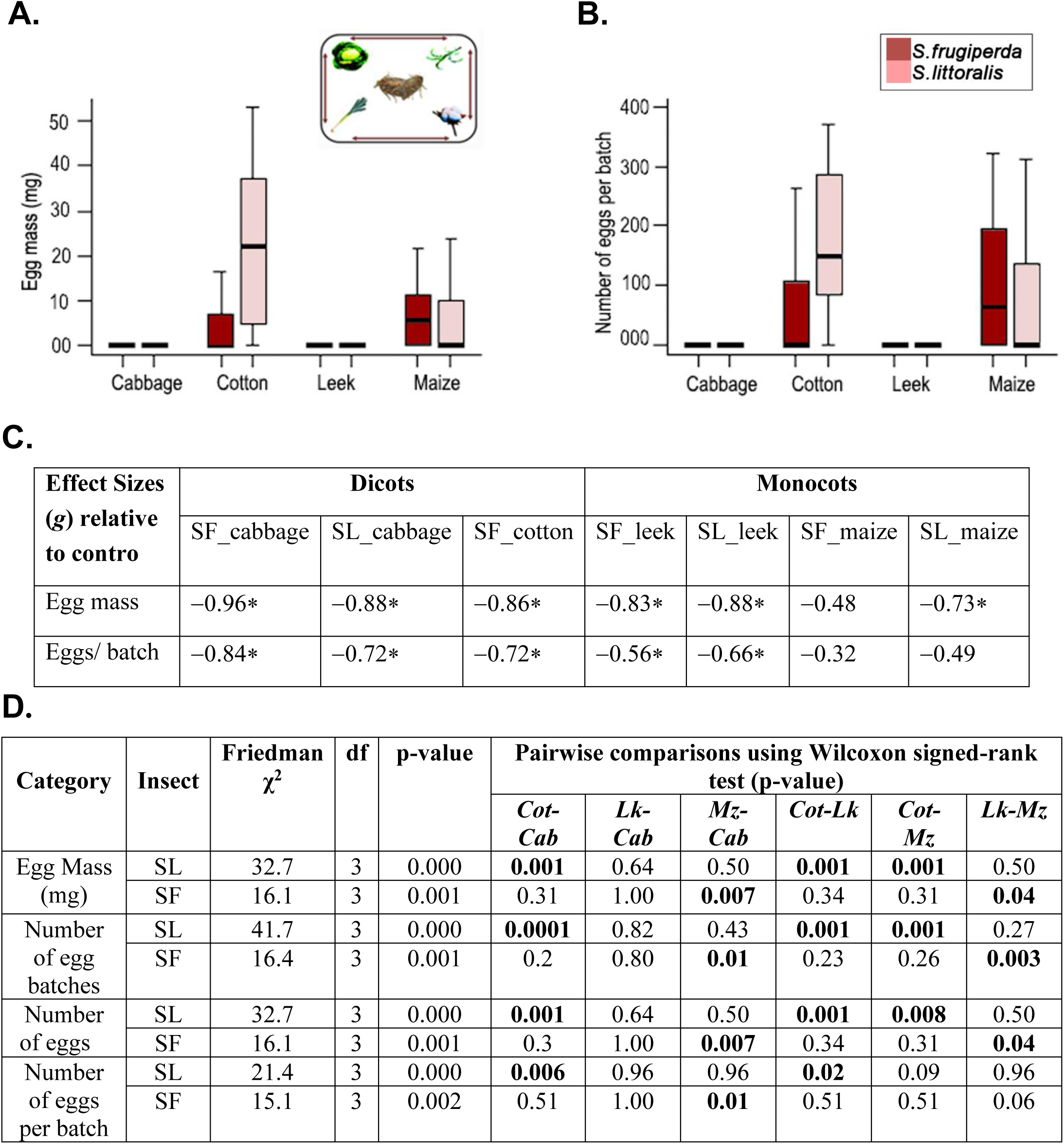
Female oviposition choice assay on the four selected host plant species. Females had the uniform chance to lay eggs on any plants under a four-choice test in bug dorms for two nights (inset). Median and interquartile ranges are plotted. Egg mass **A**) and the number of eggs per batch **B**), laid by one mated female of either *Spodoptera frugiperda* (SF) or *Spodoptera littoralis* (SL). **C**) Comparisons of means by effect size (Hedges’ *g*) against the control (SL_cotton, hence no *g* value given). **D**) Statistical analysis (all N=31) table with Friedman rank sum test followed by pairwise comparisons using Wilcoxon signed-rank test. *P*-value adjustment method: Holm-Bonferroni (Haynes 2013). The bold **p**-values indicated significant comparisons in the table. Plant acronyms: **Cab**) cabbage, **Cot**) cotton, **Lk**) leek, and **Mz**) maize.

The effects were quite strong when comparing means for the two dicot plants, in both the egg mass and numbers, for SL on cotton (as a control group) compared to SL on the cabbage and for all the SF data (**Fig. 7 C**). Correspondingly, the differences were highly significant for the SL preference to lay eggs on cotton compared to all other plant species (χ2 = 32.7, *p* < 0.001) (**Fig. 7 D**). While SF larvae performed better on leek than cotton, mated SF females’ favoured cotton over leek for oviposition. Unlike SL, the SF females showed highly significant oviposition preferences (χ2 = 16.1, p < 0.01) and preferred maize to cabbage and leek. Thus, the overall adult oviposition preferences showed a weak correlation of host performance and preference assays in the females of two tested *Spodoptera* for the four plants.

## 4 Discussion^1^

In an ecosystem, recourse utilization of herbivore insects considerably relies on dietary specialization. Dietary specialization also shapes various tritrophic interactions of herbivore insects within the ecosystem (Singer *et al*. 2014). It may also influence the correlation between female oviposition preference and offspring performance, given that the insects possess a limited neural capacity to process complex information (Egan & Funk 2005). Insects with limited diet breadth can be predicted to be adapted to make a better decision simply because they have limited resources or hosts to choose from.

The global distribution of herbivore diet breadth has been demonstrated to be continuous, with the majority specialized locally on a single plant family (Forister *et al*. 2015; Hardy *et al*. 2020). Only less than 10% of herbivore insects are true generalists thriving on more than three plant families (Bernays & Graham 1988). In this regard, both SL and SF are generalists. However, they do differ in their generalization scale. This difference in diet breadth can influence the magnitude of damage an insect pest can cause to its chances for further host range expansion and invasion of new territories.

The results of the no-choice feeding bioassay, where larvae from the neonate stage to pupation were reared on different plants, showed that SL developed and survived better on a broader range of the selected plants compared to SF. A more detailed comparison showed that SL performed better on most of the tested plants, including those they did not often encounter in their natural habitats, such as mint (*Mentha aquatica*), leek (*Allium ampeloprasum*), sunflower (*Helianthus annuus*), mad dog weed (*Alisma plantago-aquatica*), gypsywort (*Lycopus europaeus*), meadowsweet (*Filipendula ulmaria*) and blue iris (*Iris spuria*). On the contrary, SF larvae did better on two monocots, maize and leek. However, SF showed the lowest performance on other monocots, such as mad dog weed and blue iris, indicating their primary host specialization but not general monocot specialization. Our results also showed that there were also some true non-host plants in nature for both SL and SF, such as common reed, marsh marigold, common skullcap, Malabar nut, snapdragon and woody orchid.

The difference between SL and SF in development and survival on the range of plants tested was manifested by SL, e.g. showing a better overall fitness on a more comprehensive range of plants, including wetland species. A broader analysis by grouping the tested plants revealed an impact of plant systemic positions such as dicot or monocot on the feeding performance. Furthermore, the cultivation status of the host plant as a single factor considerably influenced the larval mass, but the difference between larval masses in SL and SF was independent, and no significant interaction between factors was found (**Tab. 1C; Fig. 2 B**). On the other hand, the host plant habitat (terrestrial/wetland) strongly influenced larval performance and showed significant interactions, *i.e.* different reaction norms^2^ of the two insects for the two habitat types (**Tab. 1B, Fig. 2 A**).

Larval performance in insects is often positively correlated with larval weight gain and the time to emerge as adults. The quantity of plant material consumed can be a proxy for such performance measurements (Knolhoff & Heckel 2014), as dietary consumption and energy conversion inside the gut are also positively influenced by insect survival and growth rate. However, it is worth mentioning that multiple factors can impact the amount of diet intake by an herbivore insect, such as plant nutrient content and plant defence compounds. The consumption rate on a low protein-containing plant diet may be higher than the high protein-containing diet to compensate for low nutritional value and maintain the growth rate. Similarly, a plant diet containing a high amount of plant defence is less consumed by the larvae to avoid intoxication. Also, physiological considerations such as digestibility and energy conversion efficiency of plant diets can be crucial for a better understanding of host use in polyphagous insects (Waldbauer 1968).

We observed a correlation between larval feeding performance and the nutritional indices RGR, ECI, and ECD. For example, all the above nutritional indices parameters were higher when feeding on preferred host plants than on suboptimal or non-host plants for both SL and SF larvae. Taking all measured nutritional indices into account, the dietary suitability of SL larvae for tested diets is the following: artificial diet > cotton > cabbage > leek> maize > Malabar nut. For SF, the order of diet suitability between the diets tested was artificial diet > maize > leek > cabbage > cotton > Malabar nut. Furthermore, for both insects, RGR, ECD, and ECI values for the tested host plants are close to the artificial diet, giving accurate insight into host suitability. Conversely, the wet weight gain percentage did not always correlate with host suitability or show an accurate picture. This could, for example, depend on the percentage of water content in the plant materials. Furthermore, we also found a higher mortality percentage in cotton compared to cabbage for SL, while RGR, ECI, and ECD were higher in cotton than in cabbage. Thus, the two factors showed a difference in the suitability of cotton as a host for SL larvae. As cotton is a preferred host plant for ovipositing females while cabbage is not (Thöming et al. 2013), the performance data only partly confirm the female choice of the host plant. However, the effects of host plant feeding may also have trans-generational effects. In a previous experiment, SL larvae were found to have substantially higher survival in second-generation cotton than second-generation cabbage-fed SL larvae (Houot et al. unpublished data). Altogether, comparing bioassay and nutritional indices assay results of SL and SF larvae provides insights into the correlation between insect digestive physiology and performance on diets.

Naturally, insects with more extensive diet breadth showed relatively less variation in performance upon host shifts, leading to a decoupling of performance-preference correlation in larvae (Gripenberg *et al*. 2010). Our study found mixed support for a link between preference and performance. In the larval feeding choice assay, the larval performance –preference matched well for SL. Their performance order in bioassay (Cabbage > cotton > leek > maize) matched nicely with their feeding preference. In SF, the positive correlation between larval performance and preference was limited to the most preferred plant, maize (their host). However, SF larvae performed well on leek but did not prefer to feed on that plant, suggesting their adaptive plasticity as a polyphagous pest with a precise preference for Graminid monocot plants (i.e., maize). The larval olfactory preference of both species was higher for their respective host plants, i.e., SF preferred maize over cotton, and SL preferred the opposite. However, showing no preference between cabbage and maize by both insects indicated the decoupling of association between the larval olfactory preference and performance in these two insects, which is quite common in other polyphagous insects (Gómez Jiménez *et al*. 2014).

The link between female oviposition preference, host plant rank order and larval performance rank order was weak. For example, SL larvae performed well on cabbage but did not prefer it for oviposition. Similarly, SF did not prefer leek, but SF larvae performed well on that plant. Hence, as observed in other polyphagous herbivores, no generalization can be made about larval performance and female oviposition preference in these two Spodoptera sp. (Gómez Jiménez *et al*. 2014). A similar observation was made on another polyphagous herbivore from the same genera, i.e., *Spodoptera exigua* Hübner (Berdegué, Reitz & Trumble 1998).

A higher proportion, 65%, of SL larvae fed on more than one plant compared to 37% of SF larvae, suggesting a higher degree of diet mixing nature in SL when more than one plant was available for feeding. This indicates that larval behaviour can also contribute to the higher degree of host plant use in polyphagous SL. Interestingly, plant diet mixing can improve performance in polyphagous herbivorous insects via unknown mechanisms (Unsicker *et al*. 2008; Mason & Singer 2015). According to the complementary diet hypothesis, diet mixing reduces the high amount of toxin uptake and provides superior nutrient balance to insects (Waldbauer, Cohen & Friedman 1984; Behmer, Simpson & Raubenheimer 2002). However, further dedicated studies are required to better understand such adaptive feeding behaviour in polyphagous herbivores. One factor to consider is interplant larval mobility, which is a crucial parameter to consider while studying the performance-preference correlation. SL and SF larvae showed a considerable propensity to disperse from the suboptimal host to the preferred host during the larval migration assay. In SL, laboratory and field experiments have observed directed larval migration between plants (Anderson, Sadek & Wäckers 2011; Martel *et al*. 2024).

According to the performance-preference hypothesis (alternatively, “mother knows best” hypothesis)(García-Robledo & Horvitz 2012), adult females prefer laying eggs on plants, where the larval survival is optimum (Thompson 1988), assuming that the larvae have little or no capacity to move between the plants. In such cases, adult egg-laying females have the selection pressure to choose the best host plants to maximize larval survival and fitness (Jaenike 1978). However, empirical data show a weak link between larval performance and female oviposition preference (Gripenberg *et al*. 2010; Potter, Bronstein & Davidowitz 2012) due to the contribution of other factors to the larval and adult decision-making process. Such factors can be preconditioning of a particular host plant during the larval stage (Hopkin’s principle) (Hopkins 1916) or multiple conflicting selection pressures (enemy-free space hypothesis) on female egg-laying herbivores leading to sub-optimum decision making (Ballabeni, Wlodarczyk & Rahier 2001). However, as observed in this paper, larval ability to move between plants may release the selection pressure on the females (Rivera & Burrack 2012). It is also observed that a suitable host plant choice for oviposition in terms of larval performance often depends on the diet breadth of the herbivore insect, with specialist mothers knowing better (Gripenberg *et al*. 2010). Nevertheless, female habitat preference does not inevitably correlate with larval performance (Kyogoku 2024). Hence, further evaluation of such performance–preference relations in these two generalist Spodoptera moths will be interesting.

## Conclusion

Broad host range is often considered a prerequisite for geographic range expansion and host shift events in herbivore insects (Slove & Janz 2011). The present study provides the first comprehensive data on the diet breadth of two range-expanding and highly invasive polyphagous herbivores. Our results clearly demonstrate that SL is more polyphagous than SF within the wide range of plants tested in this study. Hence, SL should have a higher potential for host range expansion, which has been observed in southern Europe, for example. However, in the last ten years, SL has established a viable population that causes extensive damage in many African countries and places in Asia. Human activities most likely assist the spread, but preferred host plants have facilitated the establishment in the new agroecosystem, e.g. maize. Our nutrition assay indicated that the larval preference for a given plant positively correlated with the larval dietary consumption and energy conversion rate in the gut, i.e. both larvae failed to utilize non-host Malabar nut plants. Apparent decoupling of larval performance and feeding preference, and female oviposition preference is observed with some minor exceptions in these two polyphagous insects, which are similar to the earlier findings on polyphagous herbivores at the species level (Gripenberg *et al*. 2010; Gómez Jiménez *et al*. 2014; Friberg, Posledovich & Wiklund 2015). Our results also demonstrated life-stage-specific host selection in tested *Spodoptera*. Interestingly, identification of life stage-specific cues may aid the development of powerful traps or lures to save plants from herbivore damage and offer an opportunity for sustainable pest management practices in countries such as Africa and southern Europe, where these insects represent a greater threat to agricultural harvests.

## Supporting information

All supplementary materials

## Authors Contributions

AR, FS, PA, DGH, ML: conceptualization; FS, PA, ML: funding acquisition; AR, SC: plant maintenance in the greenhouse; AR, NW, SC, AC, TC: conducted experiments and *Spodoptera* rearing; FS, NW, RM: statistical analysis; FS, AR, RM: Figures and tables preparation; AR: first draft preparation; PA, FS: Revised the draft; All authors have read and contributed to prepare the final draft of the manuscript.

## Acknowledgement

We want to thank SLU master student Emilia Mühlhäuser for her support while conducting some experiments. We also thank Elisabeth Marling for her technical assistance during the bioassays. Stiftelsen Olle Engkvist Byggmästare, Sweden funded this project. AR acknowledge the support by grant no. CZ.02.1.01/ 0.0/0.0/15_003/0000433, “EXTEMIT–K project,” financed by the Operational Program Research, Development and Education (OP RDE) and Excellent Team Grants (2023-2024) from Faculty of Forestry and Wood Sciences, CZU, Prague. AC is supported by grant no. CZ.02. 1.01/0.0/0.0/16_019/0000803 financed by OP RDE.

## Conflict of interest

The authors declare that the research was conducted without any commercial or financial relationships that could be construed as a potential conflict of interest.

## Footnotes

1 The corresponding data for some other dependents (time to adult, RGR1418, and pupal mass), as used in Roy et al. (2016a), are given in **Fig. Suppl. 2**, showing similar patterns to the dependents in **Fig. 1**.

2 https://en.wikipedia.org/wiki/Reaction_norm

3 https://en.wikipedia.org/wiki/Reaction_norm

## Notes

### Competing Interest Statement

The authors have declared no competing interest.

