## Supplementary material for "Diet breadth in two polyphagous *Spodoptera* moths in a wide range of host and non-host plants and the potential for range expansion": All supplementary materials

Amit Roy etal.

Supplementary Appendix 1- Classification of plants used in the current study as per Angiosperm Phylogeny group classification (APG IV)

| Scientific name (Synonym) | Common name | unranked / Subkingdo | unranked / divisio | unranked / class | Order | Family | Genus | Habitat | Cultivation |
|---|---|---|---|---|---|---|---|---|---|
| *Gossypium hirsutum* | Cotton | Angiosperm | Eudicots | Rosids/Malvids | Malvales | Malvaceae | *Gossypium* | Terrestrial | Domesticated |
| *Brassica oleracea v. capitata* | Cabbage | Angiosperm | Eudicots | Rosids/Malvids | Brassicales | Brassicaceae | *Brassica* | Terrestrial + wetland | Domesticated |
| *Sinapis arvensis L.* | Mustard | Angiosperm | Eudicots | Rosids/Malvids | Brassicales | Brassicaceae | *Sinapis* | Terrestrial | Domesticated |
| *Ricinus communis* | Castor oil plant | Angiosperm | Eudicots | Rosids/Fabids | Malpighiales | Euphorbiaceae | *Ricinus* | Terrestrial | Domesticated |
| *Trifolium alexandrium* | Egyptian clover | Angiosperm | Eudicots | Rosids/Fabids | Fabales | Fabaceae | *Trifolium* | Terrestrial | Domesticated |
| *Vigna unguiculata* subsp. *unguiculata* | Cowpea | Angiosperm | Eudicots | Rosids/Fabids | Fabales | Fabaceae | *Vigna* | Terrestrial | Domesticated |
| *Filipendula ulmaria* (L.) Maxim. | Meadowssweet | Angiosperm | Eudicots | Rosids/Fabids | Rosales | Rosaceae | *Filipendula* | Terrestrial | Wild |
| *Solanum lycopersicum* | Tomato | Angiosperm | Eudicots | Asterids/Lamiids | Solanales | Solanaceae | *Solanum* | Terrestrial | Domesticated |
| *Capsicum annuum* L. | Bell pepper | Angiosperm | Eudicots | Asterids/Lamiids | Solanales | Solanaceae | *Capsicum* | Terrestrial | Domesticated |
| *Adhota vesica (Justicia adhatoda)* | Malaber nut | Angiosperm | Eudicots | Asterids/Lamiids | Lamiales | Acanthaceae | *Justicia* | Terrestrial | Domesticated |
| *Scutellaria galericulata* | Common skullcap | Angiosperm | Eudicots | Asterids/Lamiids | Lamiales | Lamiaceae | *Scutellaria* | Terrestrial + wetland | Wild |
| *Mentha aquatica* | Mint | Angiosperm | Eudicots | Asterids/Lamiids | Lamiales | Lamiaceae | *Mentha* | Terrestrial + wetland | Wild |
| *Helianthus annuus* | Sunflower | Angiosperm | Eudicots | Asterids/Campanulids | Asterales | Asteraceae | *Helianthus* | Terrestrial | Domesticated |
| *Antirrhinum austrate* | Snapdragon | Angiosperm | Eudicots | Asterids/Lamiids | Lamiales | Plantaginaceae | *Antirrhinum* | Terrestrial | Wild |
| *Lycopus europaeus* | Gypsywort (European bugleweed) | Angiosperm | Eudicots | Asterids/Lamiids | Lamiales | Lamiaceae | *Lycopus* | wetland | Wild |
| *Beta vulgaris* | Beet | Angiosperm | Eudicots | Superasterids | Caryophyllales | Amaranthaceae | *Beta* | Terrestrial | Domesticated |
| *Caltha palustris* | Marsh marigold | Angiosperm | Eudicots | ? | Ranunculales | Ranunculaceae | *Caltha* | Terrestrial | Wild |
| *Zea mays subsp. mays* L | Maize | Angiosperm | monocot | Commelinids | Poales | Poaceae | *Zea* | Terrestrial | Domesticated |
| *Phragmites australis* | Common reed | Angiosperm | monocot | Commelinids | Poales | Poaceae | *Phragmites* | wetland | Wild |
| *Allium ampeloprasum* | Wild leek | Angiosperm | monocot | ? | Asparagales | Amaryllidaceae | *Allium* | Terrestrial | Wild |
| *Alisma plantago-aquatica* | European water-plantain (Mad dog weed) | Angiosperm | monocot | ? | Alismatales | Alismataceae | *Alisma* | wetland | Wild |
| *Iris spuria* | Blue Iris | Angiosperm | monocot | ? | Asparagales | Iridaceae | *Iris* | Terrestrial + wetland | Wild |
| *Magnolia sieboldii* K.Koch | Siebold's magnolia (Korean mountain magnolia) | Angiosperm | monocot | Magnoliids | Magnoliales | Magnoliaceae | *Magnolia* | Terrestrial | Wild |

Classification reference: An update of the Angiosperm Phylogeny Group classification for the orders and families of flowering plants: APG IV

https://doi.org/10.1111/boj.12385

? - undefined

### SUPPLEMENTAL MATERIALS Figures

#### *Supplemental Figure 1 - PCA loading plot*

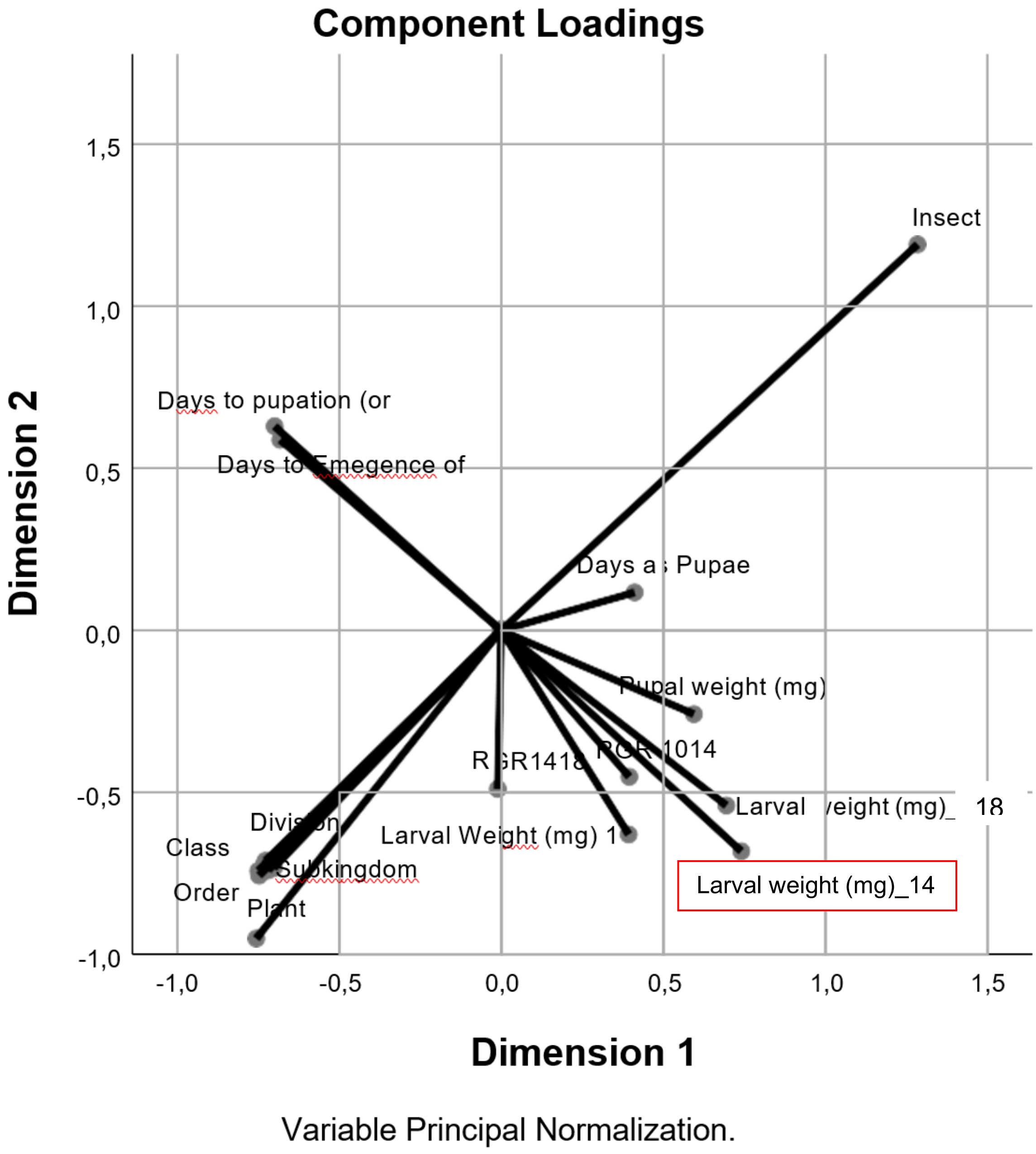

**Supplemental Figure 1.** Plot PCA-type of two dimensions showing co-relation status between the nine dependent performance variables (body masses etc) and the nine independent factors (insect, plant plus ecological and systematic levels). The dependent with highest loadings in both dimensions (**Suppl. Tab. 2**) is Larval weight (mg) at 14 days [boxed]. Several independent factors (Division, Class, Order, Family, Genus, Habitat, Cultivation) have similar values (dimensions close to –0.7) and do not separate on the plot.

#### Supplemental Figure 2 - Larval no-choice Bioassay performance additional parameters

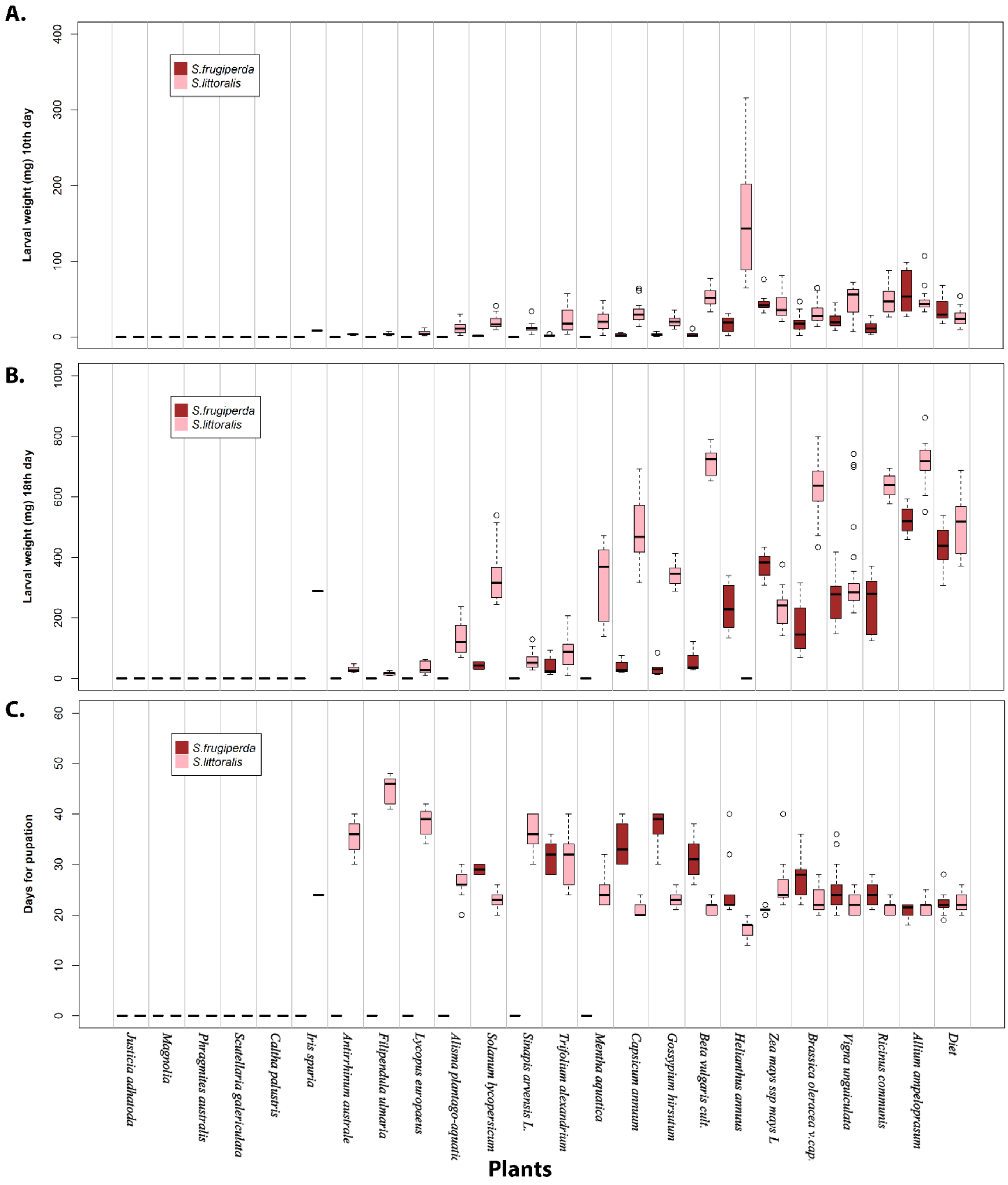

**Supplemental Figure 2.** A, B) SL and SF Larval weight after10th and 18th days, respectively, after feeding upon diverse array of host, non-host plant (23 plants) and artificial diet. Days to adult (C) same, but lower is better (higher performance), otherwise plotted to be read as in **Fig 1B&C**.

#### *Supplemental Figure 3 - Larval feeding preference behaviour in disc assay.*

**A.**

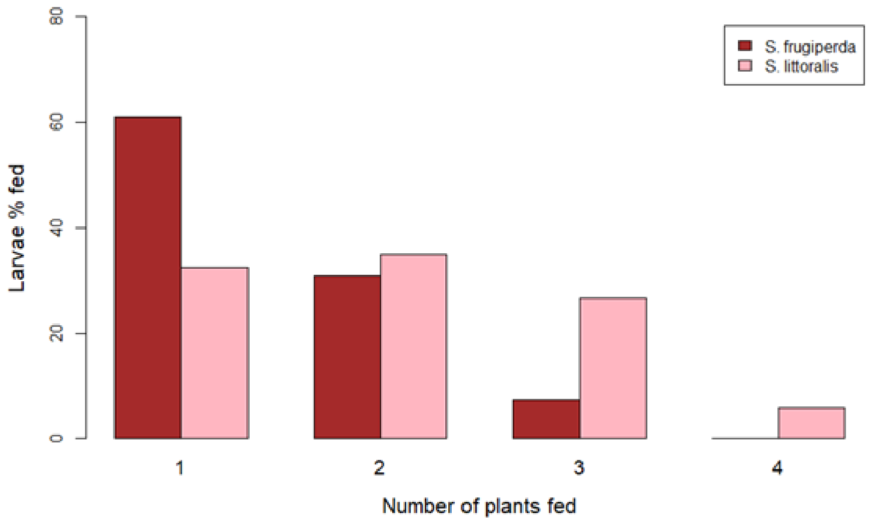

**B.**

| | Df | Deviance | Resid. Df | Resid. Dev | Pr(>$\chi^2$) |
|---|---|---|---|---|---|
| NULL | | | 7 | 205.3 | |
| Insect | 1 | 0.01 | 6 | 205.3 | 0.9405 |
| No. of Plants | 3 | 158.98 | 3 | 46.4 | < 2.2e-16 *** |
| Insect: No. of Plant | 3 | 46.35 | 0 | 0.0 | 4.771e-10 *** |

**C.**

| | ***1*** | ***2*** | ***3*** | ***4*** |
|---|---|---|---|---|
| **Difference between *S. frugiperda* and *S. littoralis*** | 1.45e-05 *** | 0.492 | 0.0001 *** | < 2.2e-16 *** |

**Supplemental Figure 1. Larval feeding behaviour in disc assay. A.** showing the percentage of SL and SF larvae feeding on 1 /2 /3 /4 plants. **B.** ANOVA table of binomial model for number of plants as factor, eaten by larvae of SL or SF. **C.** Selected contrast comparisons for number of plants eaten between SF and SL. Significance levels as in **Fig 4E**.

### SUPPLEMENTAL MATERIALS -Tables

**Correlations Transformed Variables from PCA CATREG**

| | Larval weight (mg) 10th | Larval weight (mg) 14th | Larval Weight (mg) 18d | Days to pupation (or death) | Pupal weight (mg) | Days to Emegence of Adult | Insect | Plant | RGR1014 | RGR1418 | Days as Pupae | Subkingdom | Division | Class | Order | Family | Genus | Habitat | Cultivation |
|---|---|---|---|---|---|---|---|---|---|---|---|---|---|---|---|---|---|---|---|
| Larval weight (mg) 10th[a] | 1,000 | ,769 | ,300 | –,432 | ,489 | –,443 | ,253 | ,001 | –,212 | –,111 | ,182 | –,105 | –,105 | –,091 | –,105 | –,022 | –,069 | –,106 | –,087 |
| Larval weight (mg) 14th[a] | ,769 | 1,000 | ,334 | –,563 | ,598 | –,539 | ,203 | ,003 | ,219 | –,026 | ,173 | –,122 | –,122 | –,112 | –,078 | –,021 | –,123 | –,122 | –,073 |
| Larval Weight (mg) 18d[a] | ,300 | ,334 | 1,000 | ,077 | ,440 | ,081 | ,200 | –,172 | –,053 | –,086 | ,064 | –,453 | –,453 | –,399 | –,210 | –,166 | –,256 | –,458 | –,441 |
| Days to pupation (or death)[a] | –,432 | –,563 | ,077 | 1,000 | –,277 | ,890 | ,039 | –,133 | –,354 | –,152 | –,208 | ,000 | ,000 | ,023 | ,019 | –,001 | ,061 | –,018 | –,051 |
| Pupal weight (mg)[a] | ,489 | ,598 | ,440 | –,277 | 1,000 | –,266 | ,329 | ,013 | ,137 | ,021 | ,113 | –,312 | –,312 | –,217 | –,173 | –,112 | –,219 | –,230 | –,264 |
| Days to Emegence of Adult[a] | –,443 | –,539 | ,081 | ,890 | –,266 | 1,000 | ,038 | –,144 | –,321 | –,135 | ,069 | ,018 | ,018 | ,027 | ,033 | ,005 | ,062 | –,003 | –,032 |
| Insect | ,253 | ,203 | ,200 | ,039 | ,329 | ,038 | 1,000 | –,004 | –,150 | –,083 | ,100 | ,017 | ,017 | ,007 | ,013 | ,016 | ,019 | ,002 | ,009 |
| Plant | ,001 | ,003 | –,172 | –,133 | ,013 | –,144 | –,004 | 1,000 | ,089 | ,062 | –,017 | ,181 | ,181 | ,294 | ,330 | ,352 | ,324 | ,281 | ,216 |
| RGR1014[a] | –,212 | ,219 | –,053 | –,354 | ,137 | –,321 | –,150 | ,089 | 1,000 | ,274 | –,034 | –,032 | –,032 | –,050 | ,006 | –,015 | –,093 | ,020 | ,020 |
| RGR1418[a] | –,111 | –,026 | –,086 | –,152 | ,021 | –,135 | –,083 | ,062 | ,274 | 1,000 | ,016 | –,011 | –,011 | –,025 | ,002 | ,010 | –,035 | ,006 | ,009 |
| Days as Pupae[a] | ,182 | ,173 | ,064 | –,208 | ,113 | ,069 | ,100 | –,017 | –,034 | ,016 | 1,000 | ,056 | ,056 | ,022 | ,024 | ,035 | ,037 | ,043 | ,054 |
| Subkingdom[b] | –,105 | –,122 | –,453 | ,000 | –,312 | ,018 | ,017 | ,181 | –,032 | –,011 | ,056 | 1,000 | 1,000 | ,663 | ,627 | ,607 | ,691 | ,759 | ,938 |
| Division[b] | –,105 | –,122 | –,453 | ,000 | –,312 | ,018 | ,017 | ,181 | –,032 | –,011 | ,056 | 1,000 | 1,000 | ,663 | ,627 | ,607 | ,691 | ,759 | ,938 |
| Class[b] | –,091 | –,112 | –,399 | ,023 | –,217 | ,027 | ,007 | ,294 | –,050 | –,025 | ,022 | ,663 | ,663 | 1,000 | ,389 | ,304 | ,415 | ,498 | ,561 |
| Order[b] | –,105 | –,078 | –,210 | ,019 | –,173 | ,033 | ,013 | ,330 | ,006 | ,002 | ,024 | ,627 | ,627 | ,389 | 1,000 | ,872 | ,756 | ,278 | ,567 |
| Family[b] | –,022 | –,021 | –,166 | –,001 | –,112 | ,005 | ,016 | ,352 | –,015 | ,010 | ,035 | ,607 | ,607 | ,304 | ,872 | 1,000 | ,621 | ,355 | ,607 |
| Genus[b] | –,069 | –,123 | –,256 | ,061 | –,219 | ,062 | ,019 | ,324 | –,093 | –,035 | ,037 | ,691 | ,691 | ,415 | ,756 | ,621 | 1,000 | ,481 | ,598 |
| Habitat[b] | –,106 | –,122 | –,458 | –,018 | –,230 | –,003 | ,002 | ,281 | ,020 | ,006 | ,043 | ,759 | ,759 | ,498 | ,278 | ,355 | ,481 | 1,000 | ,768 |
| Cultivation[b] | –,087 | –,073 | –,441 | –,051 | –,264 | –,032 | ,009 | ,216 | ,020 | ,009 | ,054 | ,938 | ,938 | ,561 | ,567 | ,607 | ,598 | ,768 | 1,000 |
| Dimension | 1 | 2 | 3 | 4 | 5 | 6 | 7 | 8 | 9 | 10 | 11 | | | | | | | | |
| Eigenvalue[c] | 3,347 | 2,007 | 1,141 | 1,018 | ,989 | ,779 | ,721 | ,438 | ,374 | ,120 | ,066 | | | | | | | | |

a. Missing values were imtable to textputed with the mode of the quantified variable. b. Supplementary variable. c. Eigenvalues of correlation matrix excluding supplementary variables.

#### *Supplemental Table 2 - PCA Components Loadings*

**Component Loadings**

| | Dimension | |
|---|---|---|
| | 1 | 2 |
| Larval weight (mg)_10th | .696 | –.540 |
| Larval weight (mg)_14th | .742 | –.681 |
| Larval Weight (mg) 18d | .394 | –.629 |
| Days to pupation (or death) | –.699 | .629 |
| Pupal weight (mg) | .595 | –.258 |
| Days to Emergence of Adult | –.682 | .588 |
| Insect | 1.286 | 1.190 |
| Plant | –.756 | –.950 |
| RGR1014 | .396 | –.451 |
| RGR1418 | –.010 | –.489 |
| Days as Pupae | .412 | .117 |
| Subkingdom[a] | –.730 | –.720 |
| Division[a] | –.730 | –.720 |
| Class[a] | –.750 | –.741 |
| Order[a] | –.725 | –.731 |
| Family[a] | –.702 | –.733 |
| Genus[a] | –.728 | –.710 |
| Habitat[a] | –.747 | –.756 |
| Cultivation[a] | –.720 | –.740 |

Variable Principal Normalization.

a. Supplementary variable.

#### *Supplemental Table 3 - Nutritional formula indices*

*Please see Comment*

| Abbreviation | Nutritional index | Formula | Explanations |
|---|---|---|---|
| **RCR** | Relative consumption rate | $I/(B_m * T)$ | Rate of Food consumed per mean body weight per day |
| **RGR** | Relative growth rate | $B/( B_m * T)$ | Rate of body weight increase per mean body weight per day (= RCR*ECI) |
| **AD** | Approximate digestibility | $(I\text{-}F)/I$ | Assimilated portion out of food ingested (%) |
| **ECI** | Efficiency of conversion of ingested food | $B/I$ | Gross growth efficiency (%)(=RGR/RCR) |
| **ECD** | Efficiency of conversion of digested food | $B/(I\text{-}F)$ | Net growth efficiency (%) |

$B$: dry weight gain of larva during the feeding period (mg) Adapted from Ahn et al., 2011
$T$: duration of the feeding period (days)
$I$: dry weight of food eaten (mg)
$F$: dry weight of faeces produced (mg)
$B_m$: mean dry weight of the initial and final larva during the feeding period (mg)

#### *Supplemental Table 4 - Nutrition assay: descriptive statistics* and effect sizes.

| Insect | Nutritional index | | N | | | | | | | | | | | | | | | | | | | |
|---|---|---|---|---|---|---|---|---|---|---|---|---|---|---|---|---|---|---|---|---|---|---|
| | | | | No-choice assay | | | | | | | | | | | | | | | | | | |
| | | | | Artificial Diet (Ctrl) | | | Malabar Nut | | | Cotton | | | Cabbage | | | Leek | | | Maize | | | Average |
| | | | | Mean | SD | * | Mean | SD | *g*† | Mean | SD | *g*†† | Mean | SD | *g* | Mean | SD | g | Mean | SD | *g* | |
| **SL** | Bodyweight gain | Bw | 20 | 339.3 | 111.6 | | 22.8 | 15.5 | −3.9 | 246.4 | 79.5 | −0.9 | 318.7 | 93.8 | −0.2 | 334.2 | 48.2 | −0.1 | 94.2 | 34.1 | −2.9 | −1.6 |
| **SL** | Relative growth rate | RGR | 20 | 1.6 | 0.1 | | 1.1 | 0.0 | −8.2 | 1.4 | 0.1 | −2.2 | 1.3 | 0.1 | −3.3 | 1.3 | 0.0 | −3.6 | 1.1 | 0.1 | −6.9 | −4.8 |
| **SL** | Dry weight of the Frass | F | 20 | 68.8 | 10.2 | | 25.5 | 5.0 | −5.3 | 35.9 | 10.7 | −3.1 | 36.4 | 7.5 | −3.5 | 32.7 | 4.5 | −4.5 | 17.4 | 7.8 | −5.5 | −4.4 |
| **SL** | Efficiency of conversion of Digested Food | ECD | 20 | 94.7 | 20.0 | | 28.2 | 10.8 | −4.1 | 92.1 | 26.6 | −0.1 | 77.5 | 22.5 | −0.8 | 80.5 | 15.5 | −0.8 | 43.6 | 13.1 | −3.0 | −1.7 |
| **SL** | Efficiency of conversion of Ingested Food | ECI | 20 | 47.3 | 6.8 | | 16.2 | 5.0 | −5.1 | 42.5 | 8.6 | −0.6 | 33.1 | 11.5 | −1.5 | 41.7 | 7.4 | −0.8 | 23.9 | 5.1 | −3.8 | −2.3 |
| **SL** | Relative consumption rate | RCR | 20 | 1.5 | 0.1 | | 0.9 | 0.2 | −3.5 | 1.0 | 0.2 | −2.6 | 1.0 | 0.2 | −3.1 | 1.0 | 0.1 | −3.7 | 0.7 | 0.3 | −3.2 | −3.2 |
| **SL** | Mean all indexes | | | | | | | | −5.0 | | | −1.6 | | | −2.1 | | | −2.2 | | | −4.2 | |
| | | | | | | | | | | | | | | | | | | | | | | −3.3 |
| SF | Body weight gain | Bw | 20 | 204.9 | 30.2 | | 10.3 | 11.8 | −8.3 | 111.0 | 52.4 | −2.2 | 150.9 | 38.7 | −1.5 | 184.1 | 28.8 | −0.7 | 160.3 | 37.3 | −1.3 | −2.8 |
| SF | Relative growth rate | RGR | 20 | 1.5 | 0.0 | | 1.1 | 0.1 | −8.4 | 1.3 | 0.1 | −3.2 | 1.2 | 0.1 | −3.7 | 1.4 | 0.1 | −2.2 | 1.3 | 0.1 | −3.5 | −4.2 |
| SF | Dry weight of the Frass | F | 20 | 55.0 | 12.8 | | 8.9 | 5.9 | −4.5 | 28.2 | 8.5 | −2.4 | 21.3 | 7.9 | −3.1 | 29.0 | 6.6 | −2.5 | 33.5 | 5.8 | −2.1 | −2.9 |
| SF | Efficiency of conversion of Digested Food | ECD | 20 | 98.9 | 13.9 | | 22.6 | 11.1 | −5.9 | 63.4 | 16.4 | −2.3 | 59.2 | 18.3 | −2.4 | 98.7 | 11.7 | −0.02 | 97.4 | 25.8 | −0.1 | −2.1 |
| SF | Efficiency of conversion of Ingested Food | ECI | 20 | 50.5 | 6.9 | | 14.2 | 7.9 | −4.8 | 33.3 | 9.4 | −2.1 | 34.5 | 12.0 | −1.6 | 51.9 | 7.5 | 0.2 | 41.2 | 8.7 | −1.2 | −1.9 |
| SF | Relative consumption rate | RCR | 20 | 1.3 | 0.1 | | 0.6 | 0.3 | −2.7 | 1.0 | 0.3 | −1.2 | 0.9 | 0.3 | −1.8 | 0.8 | 0.1 | −2.8 | 0.9 | 0.2 | −2.1 | −2.1 |
| SF | Mean all indexes | | | | | | | | −5.8 | | | −2.2 | | | −2.4 | | | −1.3 | | | −1.7 | |
| **SL & SF** | **Grand total** | | | | | | | | −5.4 | | | −1.9 | | | −2.2 | | | −1.8 | | | −3.0 | |
| | | | | | | | | | | | | | | | | | | | | | | −2.7 |

*) The control group for effect size, thus calculation of effect size is not relevant. †) Effect Size, as standardized bias-corrected, Hedge's *g*. ††) Bold values |*g*| >1, in other words there larger than 1 SD difference between Control and Treatment food which is a large effect. [ ] (Grey square) 95%CI for *g* includes zero (no effect).

#### Supplemental Table 5 - Nutritional assay Analytical statistics

**A.**

| Model | | Summary statistics of the model | | | | ANOVA of the model | | |
|---|---|---|---|---|---|---|---|---|
| **Response variable** | **Regression formula** | **F-statistic** | ***p*-value** | **Adjusted $R^2$** | **AIC** | **Factors** | ***p* (>F)** | |
| **Bw** | *lm(log (Bw) ~ Diet * Insect)* | 153.4 | < 2.2e-16 | 0.875 | 291.16 | *Diet* | < 2,2e-16 | *** |
| | | | | | | *Insect* | < 2,2e-16 | *** |
| | | | | | | *Diet x Insect* | 7.99E-14 | *** |
| **RGR** | *lm(log (RGR) ~ Diet * Insect)* | 83.1 | < 2.2e-16 | 0.791 | -672.07 | *Diet* | < 2.2e-16 | *** |
| | | | | | | *Insect* | 0.2083 | |
| | | | | | | *Diet x Insect* | 1.369e-13 | *** |
| **F** | *lm(log(F) ~ Diet * Insect)* | 56.8 | < 2.2e-16 | 0.720 | 183.57 | *Diet* | < 2.2e-16 | *** |
| | | | | | | *Insect* | 5.839e-10 | *** |
| | | | | | | *Diet x Insect* | < 2.2e-16 | *** |
| **ECD** | *lm(log (ECD) ~ Diet * Insect)* | 49.9 | < 2.2e-16 | 0.692 | 187.86 | *Diet* | < 2.2e-16 | *** |
| | | | | | | *Insect* | 0.7151 | |
| | | | | | | *Diet x Insect* | 2.583e-14 | *** |
| **ECI** | *lm(log (ECI) ~ Diet * Insect)* | 44.1 | < 2.2e-16 | 0.665 | 139.01 | *Diet* | < 2.2e-16 | *** |
| | | | | | | *Insect* | 0.1821 | |
| | | | | | | *Diet x Insect* | 1.826e-08 | *** |
| **RCR** | *lm (RCR ~ Diet * Insect)* | 22.8 | < 2.2e-17 | 0.500 | -72.49 | *Diet* | < 2.2e-16 | *** |
| | | | | | | *Insect* | 0.0001981 | *** |
| | | | | | | *Diet x Insect* | 1.205e-05 | *** |

**B.**

| **Comparison** | **Monocot from Dicot** | | **Cabbage vs. Cotton** | | **Leek vs. Maize** | |
|---|---|---|---|---|---|---|
| Insect | *S. frugiperda* | *S. littoralis* | *S. frugiperda* | *S. littoralis* | *S. frugiperda* | *S. littoralis* |
| Bw | 0.001 *** | <0.001 *** | 0.007 ** | 0.06 | 0.24 | <0.001 *** |
| F | 0.000 *** | <0.001 *** | 0.005 ** | 0.77 | 0.14 | <0.001 *** |
| RCR | 0.18 | 0.003 ** | 0.045 * | 0.99 | 0.22 | <0.001 *** |
| RGR | <0.001 *** | <0.001 *** | 0.39 | <0.001 *** | 0.004 ** | <0.001 *** |
| ECI | <0.001 *** | 0.03* | 0.87 | 0.004 ** | 0.014 * | <0.001 *** |
| ECD | <0.001 *** | <0.001 *** | 0.46 | 0.102 | 0.72 | <0.001 *** |

**C.**

| | **Malabar Nut** | | **Cabbage** | | **Cotton** | | **Leek** | | **Maize** | |
|---|---|---|---|---|---|---|---|---|---|---|
| Bw | *0.001* | *** | *0.000* | *** | *0.000* | *** | *0.000* | *** | *0.000* | *** |
| F | *0.000* | *** | *0.000* | *** | *0.013* | * | *0.041* | * | *0.000* | *** |
| RCR | *0.001* | *** | *0.064* | | *0.995* | | *0.001* | *** | 0.023 | * |
| RGR | *0.326* | | *0.595* | | *0.006* | ** | *0.033* | * | 0.000 | *** |
| ECI | *0.124* | | *0.721* | | *0.002* | ** | *0.000* | *** | *0.000* | *** |
| ECD | *0.098* | | *0.014* | * | *0.000* | *** | *0.000* | *** | *0.000* | *** |

**Table 6. Nutritional assay. A.** Results of ANOVA for all evaluated nutritional indices parameters of the moths. Degrees of freedom was in all cases identical: Fed on = 5; Insect = 1; Fed on: Insect = 5. Meaningful interaction between factors (insect x plant) was observed for all nutritional indices parameters. **B.** Comparison of the impact of different diets (artificial and plant) within SL and SF. Significant factor levels were compared using four contrast matrices with different sets up of the compared factor levels. Malabar Nut plant (true non-host) was significantly different from others in all comparisons. Hence omitted in the table. **C.** Comparison of the impact of different plant diets between SL and SF. Nutrition indices parameters were compared using five contrast matrices with different insects on the same plant. The *p*-value for the F-test and levels of significance: *) $p < 0.05$. **) $p < 0.01$. ***) $p < 0.001$. Empty cell (space) is the NS (non-significant) result of the test.
